# Chronically implantable µLED arrays for optogenetic cortical surface stimulation in mice

**DOI:** 10.1101/2025.02.08.637222

**Authors:** Ryan Greer, Antonin Verdier, Emma Butt, Yunzhou Cheng, Ella Callas, Niall McAlinden, Martin D. Dawson, Brice Bathellier, Keith Mathieson

**Affiliations:** University of Strathclyde, Institute of Photonics, Glasgow, G1 1RD, UK; Institut Pasteur, Institut de l’Audition, Paris, 75012, France

**Author notes:** Corresponding authors; contact at or. These authors contributed equally (Ryan Greer, Antonin Verdier). Joint senior authors (Brice Bathellier, Keith Mathieson).

**Keywords:** cortical implant, chronic implantation, optogenetics, µLED array, mouse, auditory cortex, spatiotemporal stimulation, electrophysiology, behaviour

## Abstract

Cortical implants are a proven clinical neurotechnology with the potential to transform our understanding of cognitive processes. These processes rely on complex neuronal networks that are difficult to selectively probe or stimulate. Optogenetics offers cell-type specificity, but achieving the density and coverage required for chronic, high-resolution modulation remains a challenge. Here we present a 100-element µLED array (200 µm pixel pitch, 2 × 2 mm2 footprint) coupled into a miniaturised, flexible system suitable for chronic implantation and optogenetic stimulation of the surface of the mouse cortex. The µLEDs can remain stable for over 300 hours continuous operation time *in-vivo*, allowing for months-long chronic experiments. Simultaneous electrophysiology recordings confirmed robust neuronal responses corresponding to low µLED drive currents (<5 mA), minimising thermal effects and supporting future wireless operation. The spatial resolution of neuronal responses was consistent with a simulated model of light scattering in the cortical layers, enabling device optimisation. Behavioural experiments with chronically implanted mice demonstrated robust learning during discrimination tasks using spatially distinct optogenetic stimulation patterns.

## Introduction

Cortical implant technologies have received significant attention in recent years, demonstrating their potential in clinical neuroscience. Among their many applications, the transfer of information to brain networks holds particular promise for investigating and restoring lost sensation. Optogenetics enables precise modulation of neuronal activity using light, offering cell-type specificity by employing light-sensitive proteins (opsins), which are genetically introduced into specific neuronal populations. Genetic approaches can enhance our understanding of cognitive function by improving specificity; for example, by allowing the targeted activation of excitatory or inhibitory pathways [1]. This motivates the development of optical cortical implants, tailored for optogenetic neuroscience studies in behaving animals. These devices would ideally offer broad spatial coverage, combined with millisecond-scale temporal precision, high light intensities and chronic implantability [2].

Light-emitting diode (LED) technology facilitates multi-site optogenetic stimulation across brain regions [3, 4, 5, 6, 7, 8, 9], offering the potential for minimally invasive implantation and wireless operation [8, 9]. Devices with LEDs emitting at different wavelengths could further enable the selective activation of distinct opsins, supporting advanced optogenetic experiments and the exploration of complex neuronal circuits [3, 10, 11].

The development of minimally invasive optical cortical surface stimulation methods eliminate the need for device insertion into brain matter, enabling scalable interventions for studying and modulating brain activity. The cost is a reduction in both the spatial precision, and the ability to target deeper-lying brain structures. Optical stimulation of the cortical surface is commonly achieved using external light sources, such as laser scanning systems or digital light projectors (DLPs) [12]. However, the associated optical components and electronic control systems hinder translation to experiments on freely-behaving animals and pose scalability challenges across multiple animals. In contrast, LEDs or more precisely in this case, micro-LEDs (µLEDs), can offer implantable spatiotemporal control over neuronal populations in cognitive and sensory brain areas without requiring penetration into the brain tissue [10, 13, 14, 15, 16, 17, 18, 19]. When positioned on the cortical surface, understanding the optical power delivered to the brain and the propagation of light through tissue is crucial for determining activation thresholds and spatial resolution in target cortical layers. Robust encapsulation strategies are also critical for the successful deployment of these devices, addressing challenges such as biocompatibility and long-term stability of the implanted electronic components [20]. Additionally, thermal effects must be carefully evaluated, as active devices like LEDs generate heat that could alter neuronal activity or damage brain tissue [21]. Multi-site optogenetic cortical surface stimulation requires high-density µLED arrays with spacing of a few hundred microns, covering significant tissue areas (up to 4 mm^2^) [12]. Addressing the challenges of achieving high density and scale, while demostrating robust neuronal and behavioural responses from surface-implanted µLEDs, remains an open challenge for the field.

The device presented here is a 10 x 10 array of gallium nitride (GaN)-on-sapphire µLEDs that can deliver high-intensity blue light (*λ* ≈ 450 nm) to activate Channelrhodopsin-2 (ChR2)- expressing neurons. With 200 µm pitch, and a 2 x 2 mm^2^ footprint, the µLED array enables spatiotemporal stimulation across the mouse auditory cortex. The µLED array is integrated into miniaturised opto-electronic systems for *in-vivo* experimentation in mice, including a flexible, chronically implantable system which remains stable for up to 300 hours experimental time. Transgenic mice were used to guarantee uniform expression of ChR2 across all excitatory neurons (Emx1-IRES-Cre x Ai27). Simultaneous *in-vivo* electrophysiology recordings from the auditory cortex under optogenetic stimulation by the µLED array demonstrated a robust activation threshold irradiance of between 4 - 5 mW/mm^2^ on the cortical surface. This corresponds to drive currents <5 mA with a cortical window or <3 mA for direct cortical implantation, resulting in safe temperature changes even with up to nine µLEDs active simultaneously. Optical models were used to study light propagation into brain tissue, confirming spatial resolution estimates from neuronal response analyses. These estimates place the spatial resolution (spatial extent of activation in the target regions) at ∼800 µm when a cortical window is used, or ∼600 µm when the µLED array is directly implanted. Lastly, chronically implanted µLED arrays provided spatially confined optogenetic stimulation within the auditory cortex, and we observed robust learning in behavioural discrimination experiments at similar threshold irradiance levels, but over a wider illuminated area.

## Results

### µLED array systems for spatiotemporal optogenetic stimulation of the mouse cortical surface

We developed a 100-site matrix-addressable array of GaN-on-sapphire µLEDs through microfabrication (Fig. 1a, 1b, Supplementary fig. S1, Supplementary video 1). The device features a dense array of µLEDs, arranged in a 10 x 10 grid with 200 µm spacing, enabling precise individual site or patterned illumination within a cortical region. Detailed fabrication steps are provided in the Methods; briefly, the µLEDs are 40 µm squares emitting blue light at a wavelength of approximately 450 nm, close to the peak absorption of ChR2 and a number of other common opsins. Emission is through the sapphire substrate which was thinned to 150 µm, which enhances spatial resolution as the µLEDs emit across a wide range of angles demonstrating a Lambertian profile.

**Figure 1:**
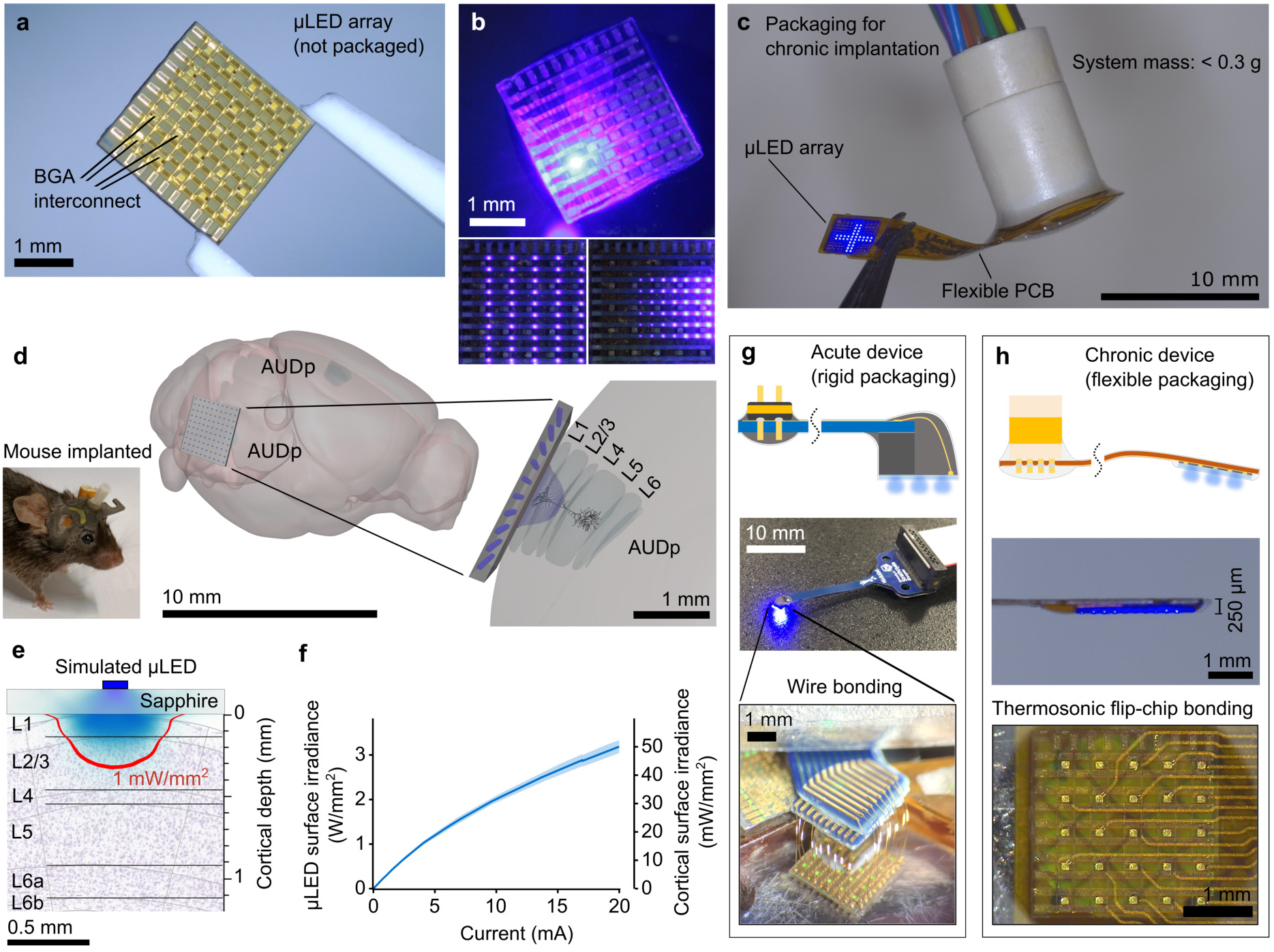
Design and modelling of µLED array systems for optogenetic stimulation of the mouse cortical surface. **(a)** Photograph of fabricated side of unpackaged µLED array, highlighting ball-grade array (BGA) interconnection pads. **(b)** Sapphire (emitting) side of µLED array showing different illumination patterns and software-adjustable brightness. **(c)** Photograph of µLED array integrated to a flexible, lightweight package for chronic implantation. **(d)** Photograph of mouse with device chronically implanted over auditory cortex (left). 3D model of µLED array over mouse auditory cortex (right); the 1 mW/mm^2^ level is shown given a single µLED illuminating a cortical surface irradiance of 10 mW/mm^2^; Illustration produced using [22, 23]. **(e)** Modelled optical profile of light propagation from a single illuminated µLED; irradiance levels are the same as (d). **(f)** Irradiance-current (L-I) characteristic at the µLED surface and cortical surface (mean and standard deviation for n = 10 µLEDs). **(g)** Schematic and photograph of rigid device for acute experimentation; zoom shows gold ball-wedge wire bonds for interconnection and µLED addressing. **(h)** Schematic of chronically implantable, flexible device and photograph of side-profile; bottom image shows thermosonic flip-chip bonding process used for highly compact interconnection.

Optogenetic stimulation of the cortical surface through an implanted µLED array enables a density of information transfer, while remaining less invasive than penetrating probes. Evaluating light propagation from the device into tissue is essential in understanding potential limitations in both the spatial resolution and the intensity of delivered light. The device was modelled using a ray tracing (Monte Carlo) simulation (Zemax OpticStudio 2021) to estimate irradiance within the cortical layers (Fig. 1d, 1e, Supplementary fig. S3a). Optical modelling indicates with a 10 mW/mm^2^ cortical surface irradiance, the 1 mW/mm^2^ contour (approximate activation threshold for ChR2) penetrates through layer 2/3 assuming the sapphire surface is implanted directly on the cortical surface (Fig. 1e). The µLEDs were optically characterised at both the µLED surface and cortical surface (Fig. 1f). For acute electrophysiology experiments, a 150 µm thin cortical window was used which increases the emitter-cortex distance, therefore increasing the illuminated area on the cortical surface, reducing the irradiance (Supplementary figs. S1d, S3b). A cortical surface irradiance of 10 mW/mm^2^ can be achieved with a drive current of less than 5 mA when the sapphire surface is in direct contact with the tissue; at this drive current range the µLED voltage is approximately 4 V (Supplementary figs. S1c, S1d).

*In-vivo* electrophysiology experiments were carried out with the µLED array placed within a compact craniotomy in an acute preparation. This necessitated a miniaturised rigid package to lower the µLED array onto a cortical window and allow space for a silicon probe to be inserted (Fig. 1g, 3a). The acute device packaging was achieved using gold wire bonds encapsulated with a potting epoxy (Supplementary fig. S2a). Chronic implantation of the µLED arrays was then made possible by optimising the packaging process through the development of a lightweight, flexible device (Fig. 1c, 1h, Supplementary fig. S2b). Highly compact interconnection was achieved using thermosonic flip-chip bonding, reducing the thickness of the device tip to 250 µm and enabling implantation to be flush with the mouse skull and surrounding tissue (Fig. 1d, Supplementary fig. S8a). The chronic device enabled behavioural experiments to be carried out through repeated optogenetic stimulation of the same neuronal ensembles within the auditory cortex, over several days. The µLED array design methodology enables the rapid development of custom packaging to facilitate a range of *in-vivo* experimental paradigms, brain regions or different animal models. Furthermore, monolithic packaging of multiple µLED arrays could enable high-density information transfer within and between cortical regions in a single chronic implantation.

### Thermal evaluation and lifetime testing of chronically implantable devices

Tissue heating is known to perturb neuronal activity, elevating or suppressing the firing activity of neurons [21, 24]. This is particularly important when using implantable µLEDs for optogenetic stimulation because they are actively powered, dissipating most of their power as heat (Supplementary figs. S4f, S4g), as well as being positioned physically close to the brain. Ideally the temperature increase would remain within the commonly quoted limit of 1 - 2 ^◦^C [25]. Our novel device geometry and a custom material stack results in complex heat flows, necessitating an in-depth thermal evaluation [26].

Infrared (IR) imaging was used in conjunction with an equivalent 2D rotationally-symmetric mathematical model replicating the flexible device, chronically implanted over cortical tissue, with appropriate material thermal properties (Supplementary fig. S4e). To verify the accuracy of the modelled device, the sapphire surface of the µLED array was imaged using an IR camera (Supplementary fig. S4a). The modelled temperature increase was compared to that measured using the IR camera under equivalent µLED power, and demonstrated good agreement (Fig. 2a). 2×2 and 3×3 squares of illuminated µLEDs were also imaged and compared with a modified thermal model; these were also found to agree well (Supplementary figs. S4j, S4k).

**Figure 2:**
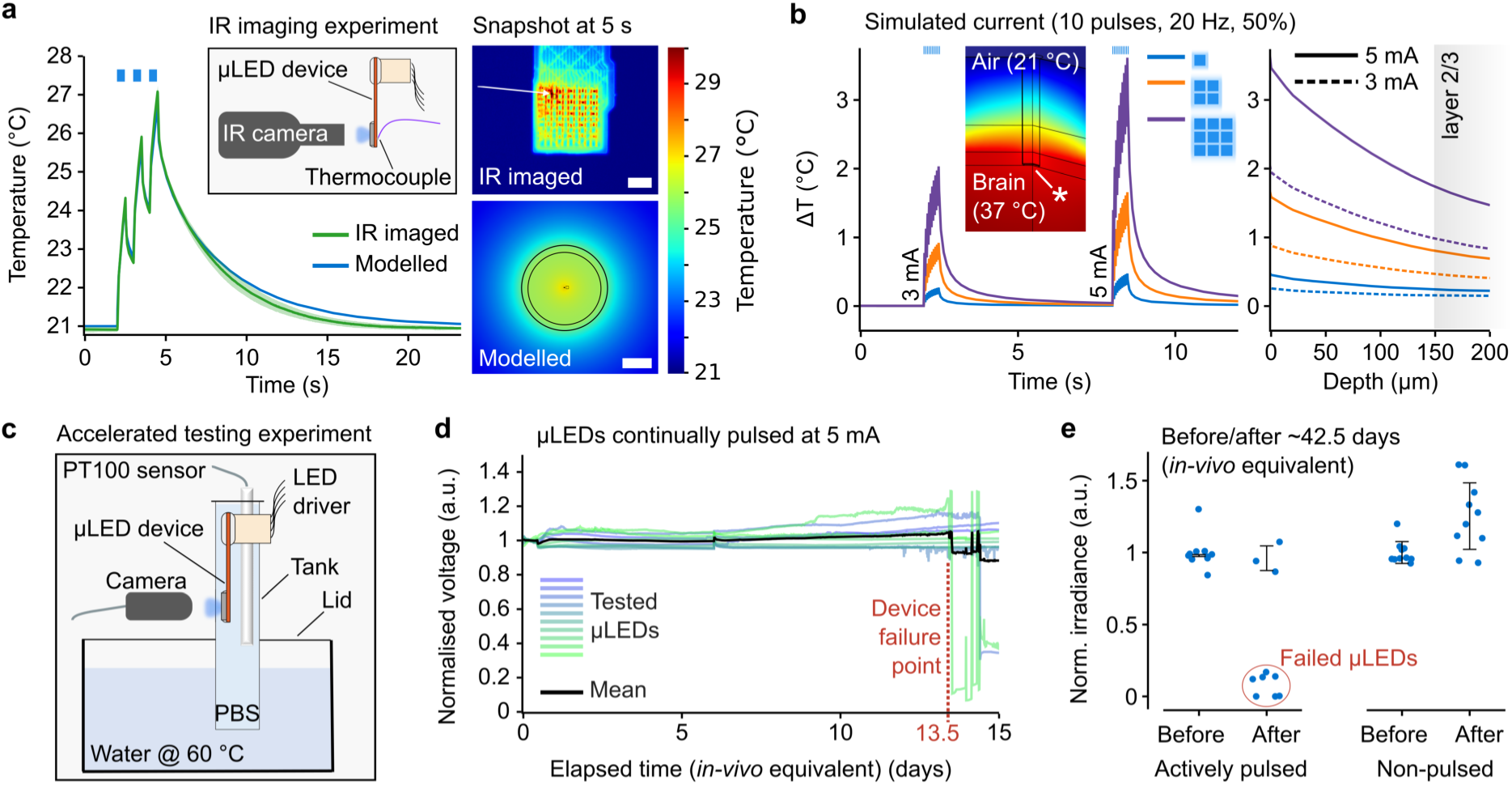
Device evaluation demonstrates minimal tissue heating and longevity of chronic operation. **(a)** Comparison of infrared (IR) imaging with equivalent thermal model (left); inset shows schematic of IR measurement setup. IR image and screenshot of modelled surface with analysed regions highlighted (right); scale bars are 1 mm. **(b)** Modelled temperature increase on brain surface (indicated by asterisk) (left); modelled heat protocol is equivalent to that used in *in-vivo* experiments, with simulated thermal power equivalent to a drive current of 3 mA and 5 mA (per µLED). Peak temperature as a function of cortical depth (right). **(c)** Schematic of accelerated ageing experiment showing the chronic device submerged in a tank of heated phosphate buffered saline (PBS) solution with µLEDs being actively driven. **(d)** Normalised measured voltage during accelerated ageing for 5 mA drive current (each µLED was activated in a repeated sequence at ∼1 minute intervals for 10 pulses at 20 Hz, 50% duty cycle); 10 tested µLEDs for a single device encapsulated with 30 µm parylene-C and medical-grade silicone. **(e)** Optical measurements of actively tested and untested µLEDs before and after accelerated ageing experiment for the same device as shown in (d).

The thermal model was then used to assess the temperature increase *in-vivo* by simulating heat pulses equivalent to 3 mA and 5 mA per µLED (Fig. 2b). The peak temperature as a function of depth is shown (Fig. 2b right). For a single µLED driven at 5 mA, the modelled peak temperature increase remained well below 1 ^◦^C. For 2×2 and 3×3 patterns, the peak temperature increase can exceed 1 ^◦^C, thus the drive current must be selected carefully. As an example, driving a 3×3 pattern at 3 mA per µLED results in a temperature increase of ∼2 ^◦^C on the brain surface and ∼1 ^◦^C in layer 2/3. Despite being minimal, temperature increases stress the need for control experiments to rule out thermal effects influencing neuronal or behavioural responses.

Accelerated ageing tests were conducted to ensure the devices would survive under the conditions of chronic implantation. Our encapsulation stack includes a thin-film oxide and SU-8 layer atop the µLEDs as well as the sapphire substrate. Subsequent packaging steps include a polyimide PCB, medical-grade epoxy underfill, parylene-C conformal coating, and a medical-grade silicone dip coating. The chronic devices were submerged in phosphate-buffered saline (PBS) solution at approximately 60 ^◦^C for two weeks (Fig. 2c, Supplementary fig. S5a). Heating of the solution allowed for an accelerated timescale to be evaluated [27]; in this case the acceleration factor was ∼4. Several µLEDs across the array were actively current-driven and the voltage recorded. The µLEDs were also monitored optically to ensure the current was driving light output and not flowing through an alternative path.

The experiment duration was over 1000 hours (42.5 days) of *in-vivo* equivalent time. Under continuous stimulation, measured voltage remained stable across all µLEDs for up to 324 hours (13.5 days) before a few µLEDs subsequently failed (Fig. 2d). To quantify the failure mechanism, µLEDs from untested rows and columns of the array were evaluated. All untested µLEDs remained functional for the entire experiment duration (Fig. 2e). The encapsulation strategy is therefore stable when the device is passive, but actively driving the µLEDs shortens the lifetime of the devices, suggesting either higher fields or thermal effects are degrading the encapsulation. These findings underline the importance of accelerated testing of active devices. Nevertheless, devices remain stable for over 300 hours of continuous operation time. Since most *in-vivo* experiments typically run for ∼3 hours per day, we conservatively estimate that the device should remain stable for at least a month of chronic experimentation. To further validate this, we evaluated devices which were chronically implanted for 29 days, actively used for behavioural experiments, and then explanted (Supplementary fig. S5c). These devices showed a minimal degradation in performance with µLEDs remaining sufficiently bright, demonstrating the robustness and re-usability of the chronically implantable devices. Furthermore, implanted devices did not have the additional medical-grade silicone layer, which has been shown to be more effective than parylene-C alone in chronically implanted devices [20].

### Simultaneous electrophysiology from auditory cortex under optogenetic stimulation with µLED array reveals activation threshold and spatial resolution

Electrophysiology recordings were conducted in the auditory cortex (AC) of awake, head-fixed mice under simultaneous stimulation from the µLED array. Recording neuronal responses to the optogenetic stimulus allowed the establishment of a threshold cortical surface irradiance for activation, and an estimation of the spatial resolution of the µLED array. Experiments were conducted in 4 transgenic mice expressing ChR2 in all excitatory neurons (Emx1-IRES-Cre x Ai27) [28, 29] (denoted ChR2+). An additional control experiment was conducted in an Emx1-IRES-Cre transgenic mouse that did not express ChR2 (denoted ChR2-). Using the µLED array, we delivered high-intensity blue light to the AC to optogenetically stimulate neuronal populations within layer 2/3. This study of irradiance thresholds and spatial resolution across distinct cortical layers provides critical insights into the precision and limitations of minimally-invasive, implantable optical methods for neuronal modulation.

For each mouse, an acute preparation consisting of a 5 mm diameter craniotomy, exposed the AC and allowed a 150 µm transparent cortical window to be implanted. A µLED array, in the acute device configuration, was lowered onto the cortical window. A 64-channel linear electrode array was then inserted through a small hole in the cortical window (0.5 mm diameter) at an angle of 28° to the cortical surface, with recording sites extending from the cortical surface to a depth of approximately 680 µm, consistent with layer 5 (Fig. 3a, 3b). Six recording sessions were conducted in total, five in ChR2-expressing mice (one mouse was repeated) and a control mouse not expressing ChR2; the control ensured that detected responses in the electrophysiology recordings were the result of optogenetic stimulation, and not thermally induced neuronal alteration of the firing rate, or stimulation artifacts. The linear electrode array was positioned at approximately the same location relative to the µLED array for all experiments (Supplementary fig. S7a). Each experiment comprised a set of trials, each trial illuminating an individual µLED at a set drive current for 10 pulses (20 Hz, 50% duty cycle) with 500 ms wait between trials. The µLED location was varied in a pseudo-random sequence for each trial. Trials were repeated up to 50 times for each combination of µLED and cortical surface irradiance. For each µLED, the drive current was individually calibrated in software to account for variations in optical output due to device fabrication. An example of optical variability of µLEDs across a device before and after the calibration is shown in Supplementary fig. S3c. The optical power of each µLED on the tested device was measured before and after the experiment. The quoted standard deviations represent the variation in cortical irradiance delivered by each µLED at the drive currents used during the experiments. This variation arises from minor degradation in the optical performance of the µLEDs, and was assumed to be due to conductive solution ingress in the acute device. As discussed, chronically implantable devices with improved encapsulation have been evaluated in solution and *in-vivo* over longer timescales.

**Figure 3:**
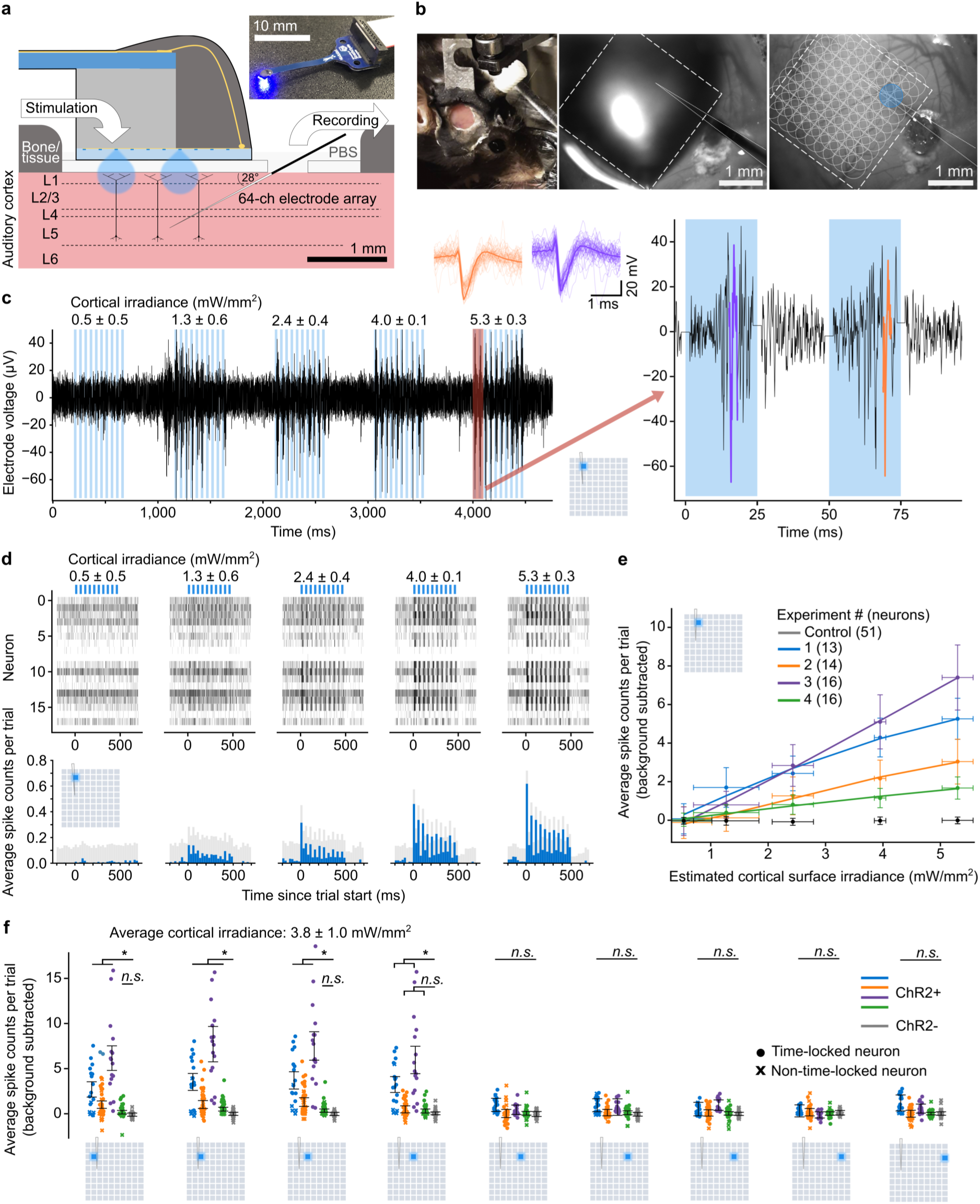
Optogenetic stimulation by µLED array elicits reliable neuronal responses from mouse auditory cortex (AC) at low drive currents. **(a)** Schematic of head-fixed *in-vivo* electrophysiology experiment. Inset shows acute device used to place µLED array over cortical window. **(b)** Photograph of Emx1-IRES-Cre x Ai27 mouse with cranial window implanted over right AC (left). Images of craniotomy with µLED array and recording probe (middle). µLED layout superimposed with an example illuminated spot (right). **(c)** Example of electrophysiology recording from a single electrode (filtered using bandpass FIR filter (cutoffs 300 Hz and 5.5 kHz), common median reference subtracted, and stimulation artefact blanked using rolling average). Each µLED is pulsed 10 times at 20 Hz, 50% duty cycle for 5 cortical irradiances (mean ± standard deviation) with corresponding µLED drive currents of 2.4, 4.0, 5.8, 8.0, 10.4 mA. Average waveforms of highlighted sorted spikes are shown (right). **(d)** Raster plot of sorted spike times under stimulation from same µLED as (c), accumulated across 50 trials (top). Peri-stimulus time histogram (PSTH) plot showing binned average spike counts per trial with mean of background subtracted, averaged over all neurons. **(e)** Dose response curves for four mice plus control, showing average spike counts per trial with background subtracted under stimulation from same µLED as (c) and (d) (ChR2+: n = 13, n = 14, n = 16, n = 16 time-locked neurons; ChR2-: n = 51 neurons. **(f)** Box plots showing average spike counts per trial with background subtracted, under stimulation from different µLEDs (ChR2+: n = 18, n = 45, n = 16, n = 53, ChR2-: n = 51 neurons); time-locked neurons are shown as circles, non-time-locked as x’s; cortical irradiances given as pooled mean ± pooled standard deviation with average µLED drive current: 8.9 ± 1.9 mA.

The following analysis identifies a threshold cortical surface irradiance needed to reliably elicit neuronal responses, which is important for determining µLED drive currents. This, in turn, determines the electronic driver specifications, has implications on possible thermal effects, and highlights key considerations for optimising optogenetic stimulation parameters. An example of a recorded waveform is shown (Fig. 3c) for 5 trials of stimulation by a single µLED, stepping through cortical surface irradiances; this µLED was chosen as it provided reasonably consistent irradiance and was located near the insertion point of the recording electrode array. Spike sorting was used to attribute the recorded electrophysiology signal to individual neurons (Fig. 3c right); only single unit activity is considered for this analysis. Spike times are shown as a raster plot for all neurons (Fig. 3d top). These were summed in 25 ms bins and averaged over all neurons to yield a peri-stimulus time histogram (PSTH) (Fig. 3d bottom). The background was taken as the mean spike counts per bin in both 200 ms periods before the 10 pulses start and after the 10 pulses end. The PSTH shows the neuronal response is time-locked to the stimulus, and is particularly strong at cortical surface irradiance values exceeding 4 mW/mm^2^. A dose response was plotted, showing average spike counts per trial with background subtracted (ChR2+: time-locked neurons, ChR2-: all neurons), against estimated cortical surface irradiance (Fig. 3e); we quote the cortical surface irradiance as an estimate due to the reduction in optical performance, as shown by the horizontal error bars. Neurons were determined as time-locked if the average spike counts per trial during the *on* period were greater than 2*σ* from the average background spike counts. Here, threshold activation is defined as a 50% success rate – 1 spike per 2 pulses on average per neuron. This is equivalent to 1 spike per 50 ms time window, which remains consistent with the off kinetics of ChR2/H134R [30]. This spike rate was achieved at a cortical surface irradiance between 4.0 ± 0.1 mW/mm^2^ and 5.3 ± 0.3 mW/mm^2^, corresponding to a drive current of between 8.0 mA and 10.4 mA. However, this is with the aforementioned reduction in optical performance observed with this device. We can expect at peak µLED performance, 5 mW/mm^2^ can be achieved at ∼5 mA (Supplementary fig. S1d). Single µLED illumination at 5 mA results in a peak temperature increase well below 1 ^◦^C on the brain surface (Fig. 2b). Neuronal responses can still be observed at lower cortical surface irradiances, suggesting an even smaller drive current is sufficient for reliable optogenetic activation. The average spike counts per trial were plotted for different µLEDs illuminated at approximately the same threshold irradiance, across the width of the device (Fig. 3f). For ChR2+ mice, mean spike counts were higher for µLEDs located near the electrode array and the control showed no time-locked response. Inter-experiment variance in the spike rate could be the result of variation in the flatness of the µLED array placement on the cortical window, or the relative positioning of the recording electrode to the µLED array (Supplementary fig. S7a). This analysis confirms that neuronal responses from ChR2+ mice are consistent with moderate cortical surface irradiances, achievable at low drive currents, and that the activation is spatially confined.

Simultaneous electrophysiology recordings under stimulation from individual µLEDs allowed examination of the spatial resolution of the device. Spatial maps were plotted showing average spike counts per trial with background subtracted for all tested µLEDs (Fig. 4a), plotted for the two experiments with the strongest responses; Experiment 1 and Experiment 3. Weighted Gaussian curves were fitted to rows 3 and 4 of the map corresponding to the highest cortical surface irradiance (Fig. 4b). The spatial resolution was inferred from the full width at half maximum (FWHM) of these fits.

**Figure 4:**
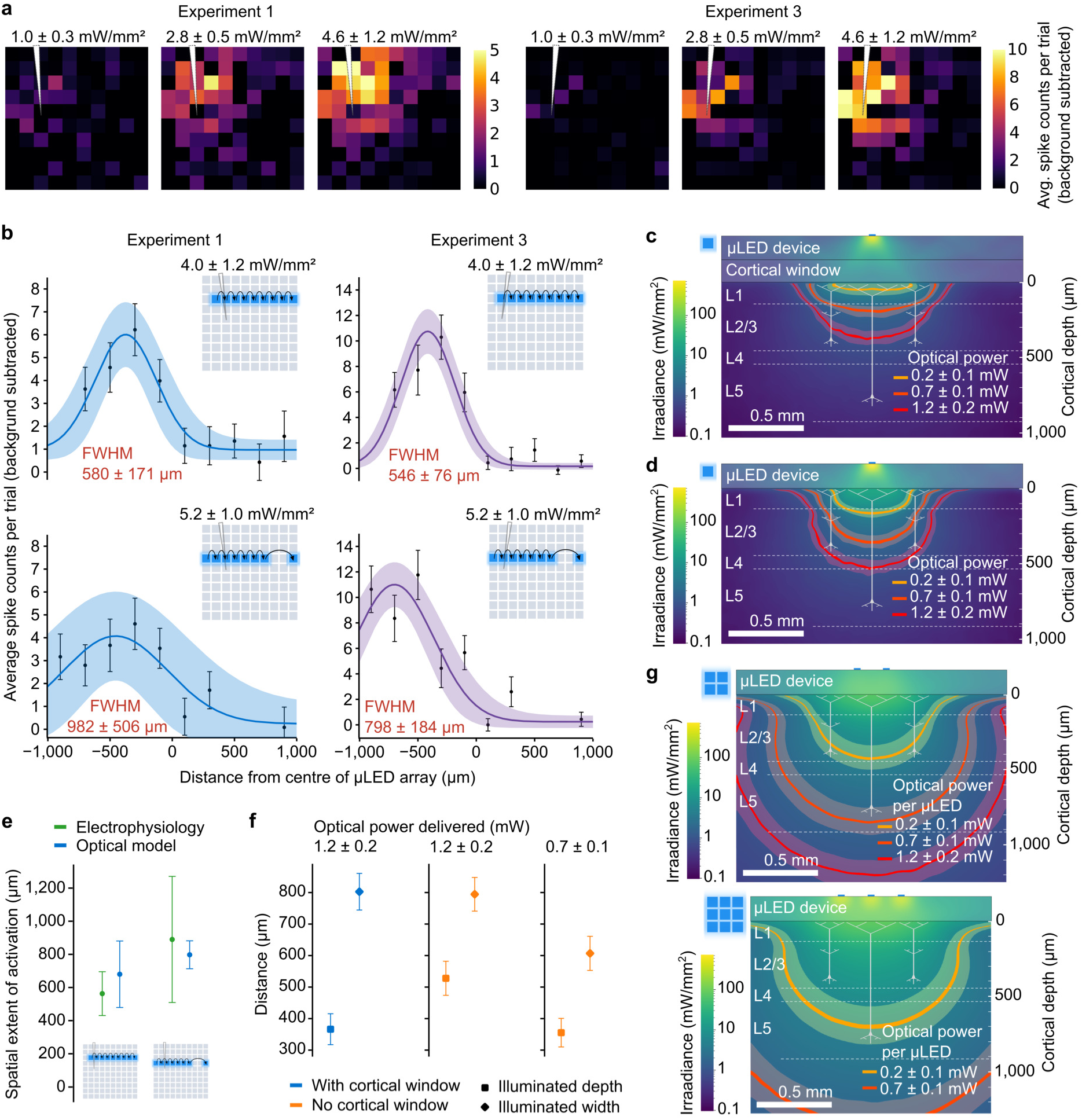
Spatial resolution identified from neuronal responses supports optical modelling. **(a)** Average spike counts per trial with background subtracted for all tested µLEDs; mean of n = 13, n = 16 time-locked neurons; cortical surface irradiances given as pooled mean ± pooled standard deviation with corresponding average µLED drive currents of 2.4 ± 0.2, 5.8 ± 0.5, 10.0 ± 0.8 mA. **(b)** Gaussian curves fitted to neuronal response to µLEDs illuminated across individual rows. Cortical surface irradiances given as pooled mean ± pooled standard deviation with corresponding average µLED drive currents of 9.7 ± 0.7, 9.8 ± 1.3 mA for each row, respectively. **(c)** Cross-sectional view of results of Monte Carlo simulation showing light propagation from µLED into brain tissue; contours show the 1 mW/mm^2^ threshold for cortical surface irradiances of 1.0 ± 0.3, 2.9 ± 0.5, 5.3 ± 0.7 mW/mm^2^. Equivalent optical power delivered into the brain tissue was determined from optical modelling. **(d)** Same as (c) but cortical window is removed and µLED array modelled directly on brain surface; cortical surface irradiances of 4.0 ± 1.1, 11.3 ± 2.0, 20.4 ± 2.8 mW/mm^2^ with corresponding optical power (and hence drive current) same as (c). **(e)** Comparison of spatial resolutions determined from the FWHM of Gaussian fits in (b) and optical model in (c); spatial extent of activation determined from the width of the 1 mW/mm^2^ threshold contour at a depth of 150 µm. **(f)** Modelled comparison of spatial resolution and depth of penetration of 1 mW/mm^2^ contour with and without a cortical window. **(g)** Summation of Monte Carlo simulation results for multiple simultaneously illuminated µLEDs, delivering the same optical power (and hence drive current) per µLED as in (c) and (d).

As described previously, optical models were developed to assess how blue light propagates from the µLED array and into the cortical tissue. To make a qualitative comparison of the optical models with the neuronal data, the 1 mW/mm^2^ contour (approximate threshold for ChR2) was plotted, given the same cortical surface irradiances evaluated during the *in-vivo* experiments (Fig. 4c). To make a comparison of the experimental spatial resolution (Fig. 4b) and the optical model (Fig. 4c), we take a cut of the model at a depth of 150 µm (approximate boundary between layer 1 and layer 2/3) and use the width of the 1 mW/mm^2^ threshold contour as an estimate of the width of activation. Experimental and modelled estimations of the width of activation are in reasonable agreement (Fig. 4e). This comparison is an approximation and implies that the optical models could be used to inform device optimisations to further improve spatial resolution. The width of activation is relatively large compared to the spacing of the stimulation sites. However, these measurements assume the highest tested cortical surface irradiance; responses can be seen at lower irradiances and therefore smaller illuminated volume and width. This comparison is also based on a cortical window between the sapphire surface and brain, which exacerbates the spread of light caused by the Lambertian emission profile from the µLEDs. Removal of the cortical window and implantation of the µLED array directly on the cortical surface is the most obvious way to improve the spatial resolution.

The 1 mW/mm^2^ contour was plotted with the µLED array modelled directly on the brain tissue (Fig. 4d). In both simulations shown in Fig. 4c and Fig. 4d, the same optical power was delivered to the brain (hence requiring the same drive current); although the cortical surface without the window is higher because the illuminated spot size is smaller (Supplementary figs. S1d, S3b). Both the depth and width of the 1 mW/mm^2^ threshold contour were compared in each case (Fig. 4f). For equivalent optical power, removal of the cortical window allows deeper penetration of light with the same lateral width (Fig. 4f middle column), while reducing the optical power to 0.7 mW results in a reduction in light spread with the same depth of penetration (Fig. 4f right column). Different patterns were also modelled based on the same delivered optical power per µLED (Fig. 4g). The activated volume significantly increases in both cases, demonstrating the ability of our µLED array to activate an entire cortical region if necessary, and at modest drive currents (Fig, 1f, Supplementary fig. S1d).

### Behavioural discrimination of spatially confined illumination patterns on auditory cortex demonstrated with chronically implanted µLED arrays

Chronic implantation of µLED arrays can facilitate repeated optogenetic stimulation of the same ensemble of neurons over days or weeks. Furthermore, the high density of stimulation sites allows activation of multiple different ensembles within the same cortical region. We trained mice to respond to artificial perceptions generated by illuminating spatially confined patterns on the surface of the auditory cortex (AC). To motivate the mice to learn the task we delivered bi-phasic electrical charge through intra-cranial stimulation of the medial forebrain bundle (MFB). This reliably activates reward centres in the brain, and the animals do not become sated as with water or food rewards [31]. Mice were first implanted with the MFB stimulator electrode, then a square-shaped craniotomy was performed to place the µLED array in direct contact with the cortical tissue after careful removal of the dura mater (Fig. 5a, Supplementary fig. S8a). The flexible packaging enabled swift and straightforward implantation, with dental cement used to secure the device to the skull. Post-surgery, mice were allowed to recover for a period of 5 days. Implanted mice were head-fixed and placed in a contention tube, with a lick detector within reach (Fig. 5b, Supplementary fig. S8b). Between behavioural experimental sessions, mice were able to remain non-encumbered due to the lightweight device packaging (Supplementary video 2), however, they were sometimes housed individually to avoid damage to the connector assembly.

**Figure 5:**
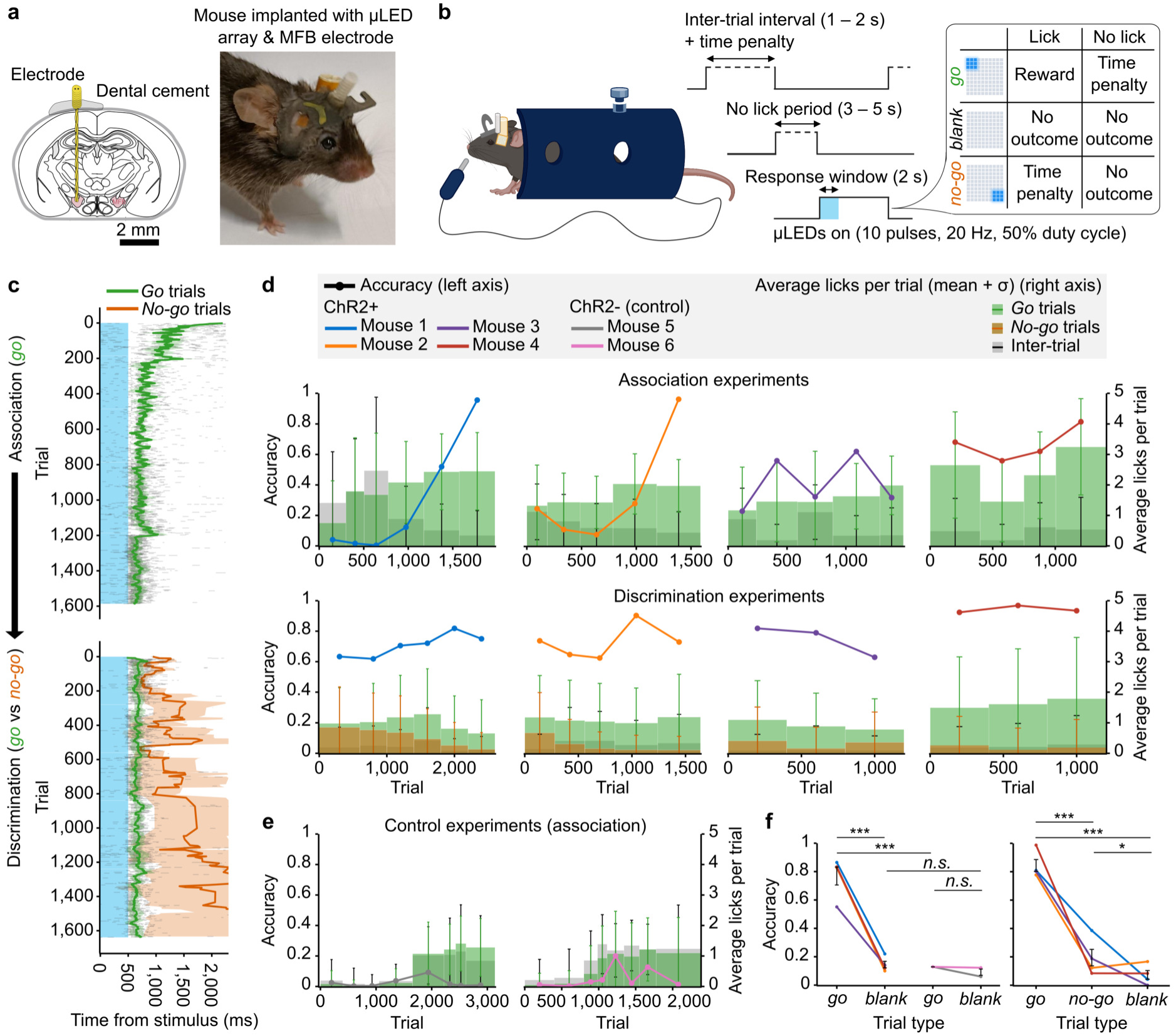
Mice chronically implanted with µLED array reliably discriminate spatially confined illumination patterns on the auditory cortex (AC). **(a)** Coronal schematic of mouse brain showing medial forebrain bundle (MFB) stimulator location (left). Photograph of mouse chronically implanted with µLED array and MFB stimulator (right). **(b)** Schematic summary of behavioural experiment; each trial consists of a period where the mouse must not lick before the optogenetic stimulus is delivered. For *go* trials, the mouse should lick the detector, and for *no-go* trials, it should not; a small number of *blank* trials providing no optogenetic stimulus are included. **(c)** Example lick plots for *Mouse 2* with mean and standard deviation shown for the lick times for the *go* and *no-go* trials. For all ChR2+ mice, the association experiment (*go* only) is followed by the discrimination experiment (*go* vs *no-go*). Control (ChR2-) mice only do the association experiment. **(d)** Learning curves for association (top) and discrimination (bottom) experiments. Each bar represents the average lick counts per trial over a session/day, with bar colour representing trial type (right axis). The thick line shows the accuracy (% of trials correct) for each session (left axis). **(e)** Same as (d) but for association experiment with ChR2- mice. **(f)** Overall accuracy for each trial type (within the best 500 consecutive trial range); association experiment (left), discrimination experiment (right).

The mice are trained on two distinct tasks - the association experiment only presents *go* trials and the discrimination experiment presents both *go* and *no-go* trials; in both cases there are a small number of *blank* trials where no µLEDs are illuminated. Initially, the mouse is trained to lick in response to the *go* stimulus in the association experiment (Fig. 5c top, 5d top). After learning the association task with sufficient accuracy (usually 4-6 sessions), the mouse moves onto the discrimination experiment, where the *no-go* stimulus is introduced (Fig. 5c bottom, 5d bottom). In the discrimination experiment, the mouse should continue licking in response to the *go* stimulus while ignoring the *no-go* stimulus. Experiments were carried out over a number of sessions (1 session per day), each consisting of several hundred trials. On each trial the µLED array delivers a 10-pulse burst (20 Hz, 50% duty cycle) of illumination at between 5 - 10 mW/mm^2^ per µLED to the surface of the AC (Supplementary tables 1 and 2). A 3×3 square of µLEDs is illuminated on one corner of the diagonal depending on the trial type (*go* or *no-go*); this broad illumination area was chosen based on previous experiments using a digital light projector (DLP) system [32, 12]. The response window opens for 2 seconds on the onset of the illumination, during which the mouse has the opportunity to receive a reward of MFB stimulation if it licks in response to the *go* stimulus. If the mouse fails to lick in response to the *go* stimulus or does lick in response to the *no-go* stimulus, a time penalty is incurred which is added to the random inter-trial interval to delay the beginning of the next trial. All other responses result in no outcome, including those during the *blank* trial where no illumination is presented. Prior to the illumination starting and response window opening, the mouse must not lick the detector for a set amount of time, to avoid continuous licking to receive the reward without learning the task; an experimental trial is illustrated in Fig. 5b. Fig. 5c shows a lick raster plot for both association and discrimination experiments, with the mean and standard deviation given for the *go* and *no-go* licks within the 2000 ms response window; these are shown for the same animal (Mouse 2) which appears to learn the task well.

Learning curves were plotted to show how the accuracy evolves over the duration of the experiments (Fig. 5d). The accuracy was computed for each trial from the percentage of *go* trials where the mouse licked (for the association experiment) or the combined percentage of *go* trials where the mouse licked and *no-go* trials where the mouse did not lick (for the discrimination experiment). A trial is only counted towards the accuracy if the number of licks within the response window is greater than 2*σ* from the mean of the background lick rate. This background lick rate was determined from the average lick counts per session from the inter-trial interval over the same time range as the response window, as shown by the grey bars. Since the mice appear to stop licking once they receive a reward (Fig. 5c), this background lick rate is not necessarily present over the course of the response window, and may provide an underestimate of the accuracy. Mice consistently learned the association task, with the exception of Mouse 3 which demonstrates a higher variability in accuracy between sessions. Good accuracy is maintained during the discrimination task with all mice able to discriminate the two optogenetic stimuli, demonstrating adaptive behaviour based on the same intensity but different spatial location of the illumination pattern. Lick raster plots have been generated for each trial type and all experiments, which demonstrate elevated licking timed with the onset of the *go* optogenetic stimulus (Fig. 5c, Supplementary fig. S9). Two control mice that were not crossed with the optogenetic strain (denoted ChR2-) received the same implantation of MFB stimulator and µLED array and were trained only on the association task (Fig. 5e, Supplementary fig. S9b). In both cases, onset of licking to receive a reward was delayed until the MFB voltage was increased to encourage some licking (at approximately 1000 trials in each case). Mice began to lick continually but this was not elevated above background licking during the response window, as with the opsin-expressing mice. Mouse 5 did appear to lick timed to the illumination during the final session which can be seen in the lick raster plot in Supplementary fig. S9b, although this was not statistically significant when compared with the background lick rate. We believe this delayed learning could be the result of the mouse visually perceiving the blue light. In future experiments, the transparent area above the µLED array could be covered, or a masking light used to saturate the mouses’ vision. The overall accuracy for each trial type is summarised for the 500 consecutive trials with the highest accuracy for each experiment (Fig. 5f). This overall high accuracy and behavioural performance was achieved at cortical surface irradiance levels similar to the thresholds determined from electrophysiology experimentation (5 - 10 mW/mm^2^) but over an increased illuminated area and volume as was shown by the optical modelling in Fig. 4g. Since the cortical window is not present for chronic implantation, these irradiances can be driven at a per-µLED current of less than 3 mA (Fig. 1f, Supplementary fig. S1d). Thermal analyses suggest this current level is safe, even for up to nine µLEDs illuminated simultaneously (Fig. 2b). Spatial confinement of the illumination pattern or reduced µLED power, as well as adjusting the lateral distance between discrimination sites, could provide further insights into the limitations of artificial perception from optogenetic cortical implants.

## Discussion

In this work, we present an array of 100 matrix-addressable GaN-on-sapphire µLEDs for spatiotemporal optogenetic stimulation of the mouse cortical surface. Individual site or patterned illumination of the brain surface enables information transfer within a cortical region, with a density of stimulation sites greater than or on par with existing approaches [10, 13, 14, 15, 16, 17, 18]. Our device matches the stimulation site density of other approaches offering 200 µm stimulation site spacing [15, 16]. These approaches integrate µLEDs which are released from their original substrate, offering flexible implementations and a reduced source-tissue proxmity. We fabricate the µLEDs on a thinned sapphire substrate, therefore maintaining the stress-matching to the µLED quantum well structures [6], maintaining high-intensity sources. The sapphire additionally acts as an encapsulation barrier and heat-spreader, addressing challenges with long-term stability [20] and thermal implications of implanted µLEDs [21, 26]. The µLEDs presented here are able to drive above-threshold irradiances several hundreds of microns deep, corresponding to reliable neuronal responses and behavioural outcomes, while maintaining low temperature increases at the implanted site. Furthermore, flexible, biocompatible packaging enables chronic implantation in the mouse model. The rapid scalability of our approach, along with robustly established stimulation parameters and *in-vivo* stability makes it an ideal platform for chronic studies into cortical function.

Simultaneous electrophysiology recordings from the auditory cortex of awake mice facilitated direct measurement of activation thresholds, with robust neuronal responses being observed at cortical surface irradiances less than 5 mW/mm^2^. Consistent behavioural responses were observed at comparable levels (5 - 10 mW/mm^2^). The µLEDs can drive these irradiances at less than 3 mA when the device is in direct contact with the tissue (Fig. 1f, Supplementary fig. S1d), resulting in safe temperature increases even with nine µLEDs illuminated simultaneously (Fig. 2b). With the cortical window implanted, the drive current increases to ∼5 mA to achieve robust neuronal responses, which is still safe for single µLED stimulation as utilised in the electrophysiology experiments. Acute electrophysiology experiments include a cortical window which would further act as a thermal insulator. Nevertheless, control experiments in non-opsin-expressing mice were conducted throughout to ensure measured responses, either neuronal or behavioural, were not thermally-induced. Low device operating power further opens the prospect of wireless, battery-powered operation for advanced studies into sensory and cognitive function in the cortices of freely-behaving animals [8, 9].

Examination of spatial resolution beyond the surface and into specific cortical layers provided an insight into the capabilities of implantable optogenetic technologies for the targeted transfer of information to the brain. The spatial resolution was analysed from neuronal responses under optogenetic stimulation from individual µLEDs, and was found to be ∼800 µm. This was in reasonable agreement with an equivalent optical model of light propagation in tissue, assuming the width of the 1 mW/mm^2^ contour at the layer 1-2/3 boundary as an approximation of the spatial extent of activation (Fig. 4e). Removal of the cortical window and direct implantation of the µLED array on the cortical surface can increase the depth of light penetration or improve the spatial resolution. The spatial resolution improves by ∼200 µm, to an illuminated width of ∼600 µm; this is in addition to reducing the required drive current (Fig. 4f). Using our optical models, we investigated if the spatial resolution could be improved through light-shaping. We modelled an optical interposer which could collimate the light using small holes fabricated into an otherwise opaque material (Supplementary fig. S3d). Incorporating this interposer could reduce the illuminated width to the range of 500 - 600 µm. We found that the dominant limitation in spatial resolution then becomes the scattering of light within the tissue, with reduction of the interposer hole size (and hence illuminated spot on the cortical surface) not reducing the illuminated width further. These findings should influence the design of future optogenetic cortical implants, and provide strategies for the minimally-invasive transfer of information to the brain using light.

The ability to manipulate distinct neuronal ensembles at the intra-cortical scale, while observing behavioural outputs, is key to exploring the circuit dynamics underlying perception [12]. Minimally-invasive chronic implantation of µLED arrays with our robust, biocompatible packaging facilitated behavioural experimentation in mice over several days. Mice were trained to respond to causal artificial perceptions generated by spatially confined optogenetic stimulation of a distinct sub-region of the auditory cortex, while not responding to an identical but spatially separated stimulus. Consistent and robust discrimination performance was observed across all ChR2-expressing animals at comparable cortical irradiance levels as determined through our electrophysiology experiments, as well as in a previous study using a digital light projector system [12]. Chronic implantation of the µLED arrays allowed repeated optogenetic stimulation of the same neuronal ensembles over several days, with a relatively straightforward surgical procedure. Accelerated ageing experiments confirmed device stability for >300 hours of continuous operation time, suitable for months-long chronic experimentation if necessary (Fig. 2d). Furthermore, chronically implanted and tested devices were shown to remain operational after explantation, offering the potential of device re-usability (Supplementary fig. S5c). Fully-implantable, high-density opto-electronic devices allowing optogenetic interrogation within cortical regions can allow us to explore the neuronal dynamics involved in a range of perceptual and cognitive processes. Future studies could include reducing the illuminated area or power, as well as reducing the lateral separation of the *go* and *no-go* illuminated patterns, to further explore the possibilities of optogenetic cortical implants for the targeted transfer of information to the brain.

The Emx1-IRES-Cre x Ai27 strain of mice were used across all *in-vivo* experiments. These mice express ChR2 in all excitatory neurons, most of which are pyramidal neurons and also express the opsin in their dendrites [28, 29]. We cannot say whether responses were from summed dendritic activation, direct somatic activation, broader network effects, or a combination of these. To gain further insight, we conducted a multi-unit analysis (MUA) using a thresholding approach, providing a neuronal response as a function of recording depth (Supplementary figs. S10a, S10b). These results show robust responses at the tip of the electrode array, suggesting neuronal activity was recorded as deep as layer 5. Optical modelling suggests the light did not penetrate this deep, even at the highest tested power (Fig. 4c). This indicates we may be recording responses from layer 5 pyramidal neurons being driven through activation of their dendrites in layer 2/3, or by other second-order effects. Soma-expressing opsin variants could be employed in future studies to gain a clearer understanding of this stimulation paradigm [33]. Pairing excitation of genetically targeted neurons with hybrid electrical stimulation could offer further insight into the specificity of the stimulation. For example, fast electrical stimulation is known to recruit inhibitory neuronal networks [34]. A hybrid stimulation approach could therefore offer insights into how best to control neuronal activity, with implications in clinical devices. Transformative experiments in humans have restored lost sensation through intracortical electrical stimulation [35, 36]. However stimulated patterns do not necessarily translate to the expected perception, highlighting the need for a deeper understanding of the spatiotemporal code [37, 38]. This is likely to require cell-type specificity, and a practical way to implement the stimulation, positioning optogenetic cortical implants ideally to address these challenges.

## Method

### µLED array fabrication

The µLED arrays were fabricated on a commercial InGaN on patterned sapphire wafer before being diced, individualised, and released (Supplementary fig. S1). The sapphire substrate is patterned and strain-matched to the quantum well (QW) structures in the µLEDs resulting in high-intensity µLEDs [6], and is thinned to 150 µm before processing; after thinning, the wafer is optically polished. A palladium layer (Pd) – 100nm – is deposited using an E-beam evaporator, to define the 40 µm pixel shape and also act as a spreading layer. A thin silicon dioxide (SiO2) layer patterned by a photoresist is used as an etching mask to etch the Pd layer and p-GaN layer, forming the pixel structure. Another SiO2 layer is then used to mask the n-GaN etching, which forms the pads for wire bonding or flip-chip bonding as well as the n-type mesa, with 10 rows, each with 10 pixels per array forming the 10 x 10 array. A lift-off process using a bi-layer of photoresist deposits a layer of Ti:Au (100 nm:300 nm) along the n-type mesa, the bond pads as well as a contact on top of each p-type pixel. Two insulation layers are formed over the n-metal tracks. Firstly, 800 nm of SiO2 is deposited and vias etched open using an RIE tool. A layer of SU-8 TF 6005 photoresist is spun and patterned then hard baked at a temperature of 200 ^◦^C for 2 hours. A lift-off process is then repeated on top of these insulation layers, depositing Ti:Au (100 nm:300 nm) in columns forming p-contacts to the pixels as well as further depositing metal on the wire bonding and flip-chip bonding pads. A final layer of SU-8 TF 6005 is spun on top of these tracks, vias opened above the bond pads and hard baked. A protective layer of photoresist (SPR4.5) is spun on top of the wafer and individual arrays are released using a diamond saw dicing tool.

### µLED array packaging

The µLED array has 20 pads for interconnection, connecting to the 10 sets of anodes or cathodes, and allowing all 100 µLEDs to be matrix-addressed. There is an option to interconnect through wire bonding pads (on two edges of the device) or a ball-grade array (BGA), for flip-chip bonding (Supplementary fig. S6c).

For acute, head-fixed experimentation in the mouse model, the µLED arrays were packaged into a custom printed circuit board (PCB) system (Fig. 1g, Supplementary fig. S2 left). The acute system consists of two PCBs (0.2 mm thin) approximately 3 cm in length which are glued together, with the µLED array attached to a silicon spacer block (1.25 x 1.25 x 1 mm^3^) on the end of the PCB. The purpose of the spacer block is to allow the µLED array to be placed in the cranial recess avoiding surrounding bone and tissue while remaining flat on the cortical window (Fig. 3a). A thermally conducting and electrically insulating epoxy adhesive paste (MG Chemicals 9460TC) is used to glue the components together. This delicate placing and gluing of components is achieved using a flip-chip bonder system and a programmable adhesive dispenser. A 1.27 mm pitch 20-pin (2 x 10) rectangular connector is soldered to both PCBs allowing electrical connection to the 10 p and n tracks. The µLED array is electrically connected to 75 nm thick ENIG (electroless nickel immersion gold) tracks on each PCB through 25 µm diameter gold ball-wedge wire bonds. The wire bonds are then potted and the PCB shanks glued together (MG Chemicals 9510) (Fig. 1g). The system is coated with 7 µm of parylene-C.

The µLED array was integrated to a flexible system for chronic implantation and used in behavioural experiments (Fig. 1c, 1h, Supplementary fig. S2 right). The shank is 100 µm thin and 2 mm wide, while still breaking out 20 connections and allowing every µLED to be individually addressed. The system was developed by bumping 75 µm diameter gold balls onto a commercial flexible polyimide PCB (PCBway) and attaching the µLED array through the BGA pads using thermosonic flip-chip bonding. A medical-grade epoxy (EPOTEK MED-301-2) is then used to underfill the µLED array for electrical and mechanical protection. A miniature circular Omnetics connector is soldered at the opposite end which is potted using the same epoxy. The system is coated with one or two layers of parylene-C (15 µm each) as well as a medical-grade silicone dip coating (NuSil MED-2214).

### Optical modelling and characterisation

The µLEDs were optically modelled using a custom ray tracing (Monte Carlo) model and simulation (Zemax OpticStudio 2021 in non-sequential mode). The model consists of a single 40 µm square GaN µLED containing a light source acting as the QW (quantum well) structure (Supplementary fig. S3a). The µLED has a gold contact on the back which is modelled as a mirror. The model includes a 2 mm square sapphire substrate (150 µm), parylene-C layer (7 or 15 µm), cortical window (150 µm) and auditory cortex. The auditory cortex was modelled as grey matter, with a 150 µm layer of white matter on the surface. The optical properties were adapted from water, incorporating suitable scattering coefficients to approximate the optical characteristics of the brain; white matter: scattering coefficient, *σ*_s_ = 50*mm*^−1^, absorption coefficient, *σ*_a_ = 0.14*mm*^−1^, anisotropy index, *g* = 0.78; grey matter: *σ*_s_ = 11*mm*^−1^, *σ*_a_ = 0.07*mm*^−1^, *g* = 0.88 [39, 40]. The ray tracing simulation simulates 2.10^7^ photons.

To characterise the µLED arrays, the laboratory set up consists of a source/measurement unit and multiplexer for software-programmable addressing of individual µLEDs. The optical power of all 100 µLEDs was measured under a sweep of drive currents. Using a similar optical model but based on the geometry of our laboratory characterisation setup (Supplementary fig. S3a right), the proportion of optical power from the µLED reaching the detector (*detector scale factor*, *DSF*) was estimated using a ray tracing simulation. The optical model in Supplementary fig. S3a (left) was then used to estimate the proportion of power from the µLED entering the brain tissue (*brain scale factor*, *BSF*). Dividing by the area of the illuminated spot size on the cortical surface (approximate diameter 530 µm with window or 270 µm without window, Supplementary fig. S3b) gives the cortical irradiance in mW/mm^2^. This gives a calibration between measured optical power and cortical irradiance and therefore cortical irradiance as a function of µLED drive current. The measured optical power to irradiance calibration equation is given at the cortical surface.

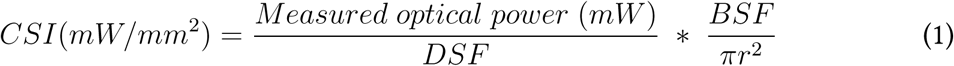

Where *CSI* is the cortical surface irradiance, *BSF* the brain scale factor, *DSF* the detector scale factor and *r* the radius of the illuminated spot on the cortical surface.

We generate regression fits for current-cortical irradiance for all 100 µLEDs in a device. The drive current can therefore be individually inferred for each µLED from the desired cortical surface irradiance during experimentation by software, allowing normalisation across the array (Supplementary fig. S3c).

### Thermal characterisation

The µLED array was imaged using an IR camera (FLIR SC7000-Series) focussed on the sapphire surface using an integration time of 1.5 ms and refresh rate of 100 Hz (Supplementary fig. S4a). Since the camera returns a digital value proportional to the number of detected IR photons, a calibration is necessary to convert this value to actual temperature in degrees celcius. The calibration uses an E-type thermocouple fixed to the back surface of the device. The sapphire surface was brought into focus first by focussing on the mesa structures in the device, and then moving the stage back through the known thickness of the sapphire (150 µm) (Supplementary fig. S4b).

An area of sapphire, located close to the µLED being tested was used for calibration (Supplementary fig. S4b). Using a heat gun, the device and thermocouple were simultaneously and uniformly heated to over 50 ^◦^C and allowed to cool down to room temperature, while both IR camera and thermocouple measurements were made at an interval of 100 ms. This process was repeated 5 times and a second-order polynomial was fitted to the thermocouple measurements against camera counts (Supplementary fig. S4c).

For the thermal measurement, the µLEDs were pulsed under several stimulation protocols. For verification of the model and to more easily assess the time constant of the thermal drop-off, a simple protocol was used; single µLED pulsed at 5 mA (1 Hz, 50% duty cycle) (Fig. 2a). The IR image in Fig. 2a shows a frame corresponding to the peak temperature from the IR camera calibrated using the thermocouple measurements. The calibration was carried out using a measurement from the sapphire area of the image (Supplementary fig. S4b) with the remaining image scaled accordingly, however this assumes linear emissivity between the materials and so is only indicative.

Thermal modelling used the COMSOL Multiphysics 5.6, heat transfer package. Two thermal models were designed; firstly to simulate the IR imaging experiment (i.e. the device suspended in air) (Supplementary fig. S4d) to verify the accurateness of the model, and secondly to simulate a chronic *in-vivo* experiment (Supplementary fig. S4e). Both are 2D rotationally-symmetric models and implement the device as a layer of gallium nitride (GaN) atop sapphire with parylene-C encapsulation. The device model also includes the polyimide PCB with a layer of copper on each side, gold bump bonds between the GaN and PCB, and an epoxy underfill. As the model is rotationally symmetric (Supplementary fig. S4i), care has been taken in designing the geometry such that the resulting 3D model has the correct volumes of each material; particularly the µLED itself which acts as the heat source in the model. The chronic implantation was simulated by placing the device over a volume of brain tissue with encapsulating cement atop, in a modelled environment (Supplementary fig. S4e). Brain tissue was approximated using the thermal properties of water.

To input thermal power to the model, firstly the electrical and optical power were measured as a function of drive current for a set of 10 representative µLEDs on one of our devices (Supplementary fig. S4f). The optical power generated by the µLED is calculated from the measured power using optical modelling as described previously. By subtracting the optical power from electrical power, thermal power can be determined (Supplementary fig. S4g). A second-order polynomial is fitted so thermal power can be calculated for any drive current and this is then input to the model as a train of pulses (3 mA/5 mA, 20 Hz, 50% duty cycle, Supplementary fig. S4h).

Since the model is rotationally-symmetric, the µLED heat source cannot be implemented as a cuboid (the true geometry of the µLED). Instead, a cylinder of equivalent volume is used and IR imaging used to verify the accurateness of this assumption. To implement multiple µLEDs at the same time, a concentric ring is used with a radius similar to the distance between the rows of µLEDs. The volume of this ring is equivalent to the volume of 3 µLEDs or 8 µLEDs, with the centre µLED having a volume equivalent to a single µLED. A similar comparison was made between IR imaging with squares of µLEDs pulsed and the model with rings, and both cases were found to agree well (Supplementary figs. S4j, S4k).

### Accelerated ageing experiments

The accelerated testing setup uses a heated water bath to ensure a steady temperature and safety for running the experiment over a long timescale. A custom-machined aluminium lid is placed over the water bath with a hole to allow a polystyrene tank to protrude through (Fig. 2c, Supplementary fig. S5a). This setup allows the device to be submerged in PBS within the tank while preventing water evaporating and escaping the chamber.

Several µLEDs across the array were pulsed constantly (approximately once each per minute) with a fixed current (5 mA, 10 pulses at 20 Hz, 50% duty cycle) and the voltage was measured to evaluate changes in the resistance of the device (Fig. 2d). A microscope camera images the device for the duration of the experiment. A PT100 sensor is also placed in the solution for accurate temperature measurements over the course of the experiment. 10 different µLEDs were tested in a repeated cycle for 337 hours (approximately 14 days), which corresponds to an accelerated time of 1025 hours (approximately 42.5 days) based on the Arrhenius relation for the accelerated testing of polymers [27].

### Animals

Optogenetic experiments were conducted on adult transgenic mice (Emx1-IRES-Cre x Ai27D) expressing ChR2 in all excitatory neurons of the pallium [29]. Control experiments were performed on adult transgenic mice (Emx1-Cre) that did not express any opsin.

### Surgery: cranial window implantation

For chronic unilateral access to the auditory cortex (AC), a cranial window was introduced and a metal post for head fixation was implanted on the opposite side. The procedure was performed on 8-12 week-old mice. Buprenorphine (Vertergesic, 0.05-0.1 mg/kg) was administered 30 minutes before anesthesia with isoflurane (3%), maintained at 1-1.5% and the animal placed on a thermal blanket during the procedure. The eyes were covered using Ocry gel (TVM Lab) and Xylocaine 20mg/ml (Aspen Pharma) was injected locally at the incision site. The right masseter was partially resected and a large craniotomy (5 mm diameter) was made above the AC, guided by skull sutures. A 5 mm circular coverslip (150 µm thick) was sealed over the craniotomy using cyanolite glue and dental cement (Ortho-Jet, Lang). For the electrophysiology experiment, a grounding electrode (PlasticsOne MS303T/2-B) was implanted contralaterally.

Post-surgery, mice received a subcutaneous injection of glucose (G30) and Metacam (1 mg/kg). The mice were then housed for one week with metacam administered by drinking water or dietgel (ClearH20) without any manipulation. All animals were housed together before and after surgery without any degradation to the implanted grounding electrode.

### Acute *in-vivo* electrophysiology experiments

Electrophysiology recordings were conducted in mice implanted with a glass cortical window above the AC for 3-4 weeks. One hour prior to *in-vivo* experiments, mice were anaesthetised with isoflurane (3% at induction), placed on a thermal blanket and maintained at 1-1.5%. Excess dental cement was trimmed to allow better access to the craniotomy. A 0.5 mm hole was carefully drilled by hand with diamond-coated drill tips in the glass window. Mice were then awakened and Kwik-Cast silicone (World Precision Instrument) was applied on the cranial window to prevent drying.

Six recording sessions were conducted in five mice: four transgenic mice expressing ChR2 (ChR2+) and one not expressing ChR2 (ChR2-) as a control. A 64-channel linear recording electrode array (NeuroNexus A1x64 Poly2-6mm-23S-160) was inserted 1.4 mm at an angle of 28^◦^ with respect to the brain surface through the previously drilled hole. The µLED array was placed on top of the cortical window above the recording electrode array (Fig. 3a, Supplementary fig. S7a). Voltage traces were recorded at 20 kHz using an RHD2000 USB interface board (Intan Technologies).

The stimulation protocol cycled through each µLED in a pseudorandom sequence, pulsing 10 times (20 Hz, 50% duty cycle) for five cortical irradiance levels: 1.0 ± 0.3, 1.2 ± 0.6, 2.8 ± 0.5, 3.2 ± 0.8, and 4.6 ± 1.2 mW/mm^2^ every 500ms. Each µLED-irradiance combination was repeated over 50, 20, or 10 trials, prioritising µLEDs near the electrode array, which was necessary due to time constraints. Eight µLEDs were tested at all five irradiances, while others were tested at three: 1.0 ± 0.3, 2.8 ± 0.5, and 4.6 ± 1.2 mW/mm^2^. Cortical irradiances quoted as pooled mean ± pooled standard deviation. Non-functional µLEDs or those with insufficient power were excluded. µLED drive currents corresponded to 2.4 ± 0.2, 4.0 ± 0.3, 5.8 ± 0.5, 7.7 ± 0.6 and 10.0 ± 0.8 mA. Currents quoted as mean ± standard deviation. Each mouse underwent a 5030-trial experiment lasting over approximately 1 hour.

### Electronic control system for *in-vivo* experimentation

A custom electronic control system was developed to power the µLED array and address all 100 µLEDs via software protocols (LabVIEW 2021, National Instruments) (Supplementary figs. S6a, S6b). The system includes an Arduino UNO R3 microcontroller for communication with the LabVIEW-equipped PC and programmable 10-channel current DACs using two DC2903A development boards (Analog Devices). A custom shield PCB connects to the Arduino UNO allowing SPI communication with the current supplies. The PCB also houses a 10-channel solidstate switch array (Texas Instruments TM7211) controlled by the microcontroller. Additionally, the microcontroller supplies a trigger signal to the Intan recording system to help align recorded data with the µLED stimulation onset and offset.

### Spike sorting and analysis of electrophysiology data

Raw electrophysiology data was automatically spike sorted using the Spike Interface package and Kilosort 2.5. Manual curation was carried out with Phy to remove incorrectly identified clusters. Stimulation artefacts were also manually identified and excluded from analysis.

A custom Python pipeline was used to aggregate spike counts based on trigger signals recorded by the Intan recording controller and to perform all subsequent data analysis. Spike counts were binned into 5 ms intervals, with each trial consisting of 100 bins (500 ms) of stimulus, 40 bins (200 ms) before stimulus onset, and 40 bins (200 ms) after stimulus offset, totaling 180 bins (900 ms). For simplicity, further binning into 25 ms bins was carried out to coincide with the stimulation protocol (20 Hz, 50% duty cycle).

The pre-stimulus and post-stimulus bins were used to calculate mean background spike counts, which were subtracted from the primary data recorded during the 500 ms stimulus. This allowed determination of which neurons were time-locked to the stimulus and the magnitude of their responses. Data visualisation was carried out using the Matplotlib and Plotly packages in Python 3.

The dose responses (Fig. 3e) show pooled mean with error bars as pooled SEM of average spike counts per trial (background subtracted); ChR2+ (optogenetic): n = 13, 14, 16, 16 time-locked neurons; ChR2- (control): n = 51 neurons. Sigmoid curves were fitted to each set of points to aid visualisation.

Boxplots (Fig. 3f) show error bars as pooled SEM of standard deviations of average spike counts per trial (background subtracted); significance levels were determined using the independent two-sample, single-tailed Student’s t-test with alt. hypothesis: spike counts (ChR2+) > spike counts (ChR2-); *p < 0.05, n.s. means alt. hypothesis rejected at p < 0.05); ChR2+ (optogenetic): n = 18, n = 45, n = 16, n = 53, ChR2- (control): n = 51 neurons.

Spatial resolution plots (Fig. 4b) show error bars as pooled SEM of standard deviations of average spike counts per trial (background subtracted); n = 13, 16 time-locked neurons. Gaussian curves were fitted to each row using data points, *x*, weighted by the error bars.

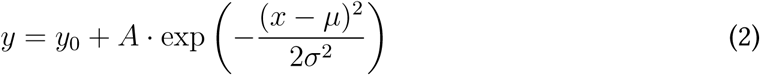

Initial values for (*y*_0_, *µ*, *A*, *σ*) were (0, −500, 50, 200) with bounds as ([0, 2], [−900, 0], [0, 200], [100, 1000]). The *x* axis is in µm. The standard error of the fitted parameters defined the shaded region. Reduced *χ*^2^ values were 0.56 and 1.40 for rows 3 and 4 of Experiment 1, and 1.68 and 4.43 for rows 3 and 4 of Experiment 3.

One experiment was omitted (Supplementary figs. S7b, S7c): a repeat on the same animal as Experiment 1, carried out a day after a recording probe broke and was manually removed. This data was considered contaminated and excluded from Figure 3.

### Surgery: chronic implantation of flexible µLED device

The procedure was performed on 8-12 week-old mice. Buprenorphine (Vertergesic, 0,05-0,1 mg/kg) was administered 30 minutes before anesthesia with isoflurane (3%), maintained at 1-1.5% and the animal placed on a thermal blanket during the procedure. The eyes were covered using Ocry gel (TVM Lab) and Xylocaine 20mg/ml (Aspen Pharma) was injected locally at the incision site. A square craniotomy of 2.4 x 2.4 mm^2^ was performed above the auditory cortex using skull bone sutures as a landmark. The µLED array was placed directly on the cortical surface after removal of the dura mater and the edges were glued to the skull using SuperGlue (Loctite). The flexible PCB packaging was fixed with dental cement (Ortho-Jet, Lang) and a MFB electrode was implanted.

Post-surgery, mice received subcutaneous glucose (G30) and Metacam (1 mg/kg). They were housed for one week with Metacam provided via drinking water or DietGel (ClearH2O) without manipulation. All animals were co-housed before and after surgery without damage to the implanted grounding electrode. However, occasional temporary isolation occurred when co-housed mice chewed on the connectors of implanted devices.

### MFB stimulation

All mice used in behavioral experiments were implanted with an intracranial electrode stereo- taxically targeting the medial forebrain bundle (MFB) to deliver a stable reward, as demonstrated previously [31].

The MFB electrode was implanted using stereotaxic coordinates (AP -1.4, ML +1.2, DV +4.8) using a NeuroStar stereotaxic device (Stereodrive 018.117). The electrode and headplate were secured to the skull using dental cement (Ortho-Jet, Lang).

MFB stimulation was provided with a pulse train generator (PulsePal V2, Sanworks) delivering 2 ms biphasic pulses for 100 ms at 50 Hz.

### Behavioural experimental procedure

The experimental setup is shown in Supplementary figs. S8b, S8c. The µLED array is controlled by the Arduino-based electronic control system, while the MFB stimulator is connected to a pulse train generator. A PC running LabVIEW (for µLEDs) and MATLAB routines manages the overall experiment. A National Instruments DAQ detects licks, applies MFB stimulation as a reward, and triggers the LabVIEW routine to apply the appropriate µLED array illumination pattern.

Mice performed behavioural tasks five days per week (Monday to Friday). Throughout the behavioural training period, food and water were provided *ad libitum*.

#### Habituation

Mice were habituated to head fixation over two days by remaining in the contention tube without reward for 20 minutes on the first day and 40 minutes on the second day. On the third day, mice received a systematic reward for licking the spout, with a maximum reward rate of 1 reward per second for 500 trials to establish the association between licking and reward. To encourage licking in non-water-deprived mice, the spout was positioned very close to their mouths. During this initial phase, before calibration, MFB stimulation was set at 3 V and reduced by the experimenter if it caused a motor reaction.

#### *Go* training (association)

Once animals were spontaneously licking, the association experiment was conducted. *Go* trials were presented with 90% probability, while the remaining trials were *blank* trials (no illumination and no reward/time-out period). A trial consisted of a random inter-trial interval between 1 and 2 seconds, a ‘no lick’ period between 3 and 5 seconds (these timings were randomised to avoid prediction of the illumination based on timing), and a fixed response window of 2 seconds. A single lick was registered as a response. Licks during the response window on a *go* trial was scored as a ‘hit’ and triggered an immediate reward via MFB stimulation. No lick was scored as a ‘miss’ and a random time-out penalty between 5 and 7 seconds added to the inter-trial interval was incurred.

#### *Go*/*no-go* training (discrimination)

Licks for *go* trials were rewarded the same as the association experiment. During presentation of the *no-go* pattern, the absence of a lick was scored as a ‘correct rejection’ and the next trial immediately followed. Any licking during the response window of the *no-go* trials was scored as a ‘false alarm’; no reward was issued and the the same random time-out penalty was incurred. Each session contained 47.5% probability of a *go* trial, 47.5% probability of a *no-go* trial and 5% probability of a *blank* trial.

All experimental parameters for the behavioural experiments are given in Supplementary tables 1 and 2.

### Behavioural data analysis

The voltage on the lick detector was recorded using a National Instruments DAQ. The timing of individual licks was determined from the activation and deactivation of the sensor (a single lick was registered when the mouse’s tongue made and then broke contact with the sensor). Licks were recorded throughout the experiment, including during the inter-trial period.

For the summary plot (Fig. 5f), error bars show SEM over mice with statistical significance levels determined using the independent two-sample, single-tailed Student’s t-test with alt. hypotheses: *go* accuracy > *no-go* or *go* accuracy > *blank* accuracy and/or ChR2+ accuracy > ChR2- accuracy. ***p < 0.001, **p < 0.01 *p < 0.05, n.s. means alt. hypothesis rejected at p < 0.05. ChR2+: n = 4 mice, ChR2-: n = 2 mice.

All relevant data are included in the article and supplementary files. Additional data requests may be directed to the corresponding author.

## Acknowledgements

The HearLight project has received funding from the European Union’s Horizon 2020 research and innovation programme under grant agreement No 964568. K.M. was supported by the Royal Academy of Engineering Chair in Emerging Technologies. The authors would also like to acknowledge the following staff within the Institute of Photonics; cleanroom support: Ronnie Roger, Beniot Guilhabert, Ian Watson; administration: Sharon Kelly; machining work: Lewis Hannah.

## Author contributions

R.G. designed and implemented the device packaging, carried out laboratory characterisation and modelling, analysed the *in-vivo* data, prepared the figures and wrote the manuscript. A.V. carried out the surgeries, *in-vivo* experiments, data collection and analysis, and co-authored the writing of the manuscript. E.B. and Y.C. designed and fabricated the µLED arrays based on an approach developed by

M.D.D. E.C. was involved in behavioural *in-vivo* experiments. N.M. assisted with laboratory experiments and device packaging. R.G., A.V., B.B. and K.M. conceptualised the devices and experimental design. All of the authors reviewed and contributed to the manuscript.

