## Supplementary figures for "Chronically implantable µLED arrays for optogenetic cortical surface stimulation in mice"

Affiliations:

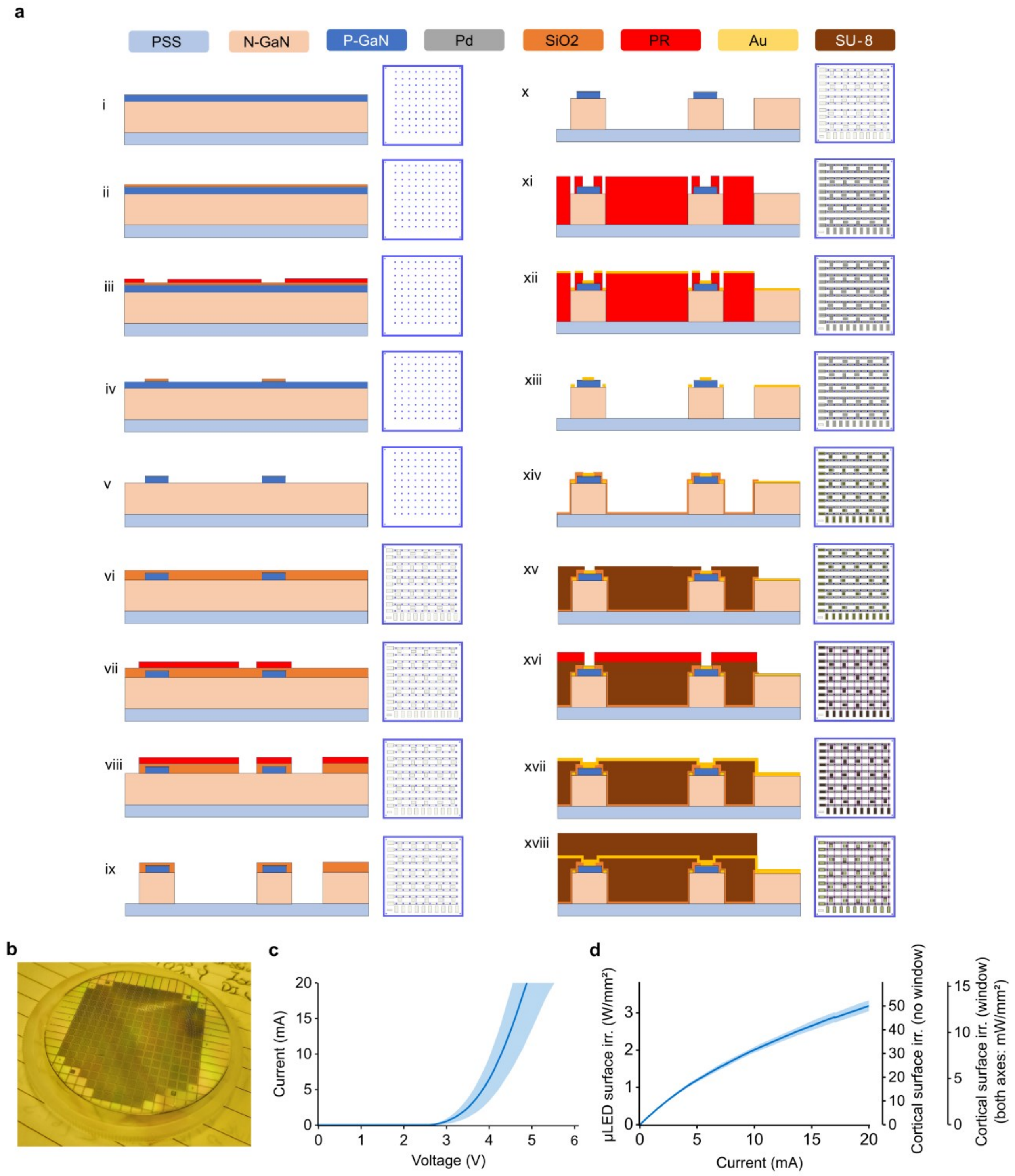


**Supplementary figure S1: Summary of µLED array fabrication process and characterisation. (a)** Cross-sectional view of fabrication steps (left), corresponding photolithography mask for each fabrication step (right). **(b)** 2-inch wafer-scale fabrication with 220 µLED arrays. **(c)** Typical current-voltage (I-V) characteristic. **(d)** Typical irradiance-current (L-I) characteristic, including at the cortical surface with and without a cortical window present. Both (c) and (d) show mean and standard deviation of n = 10 µLEDs.

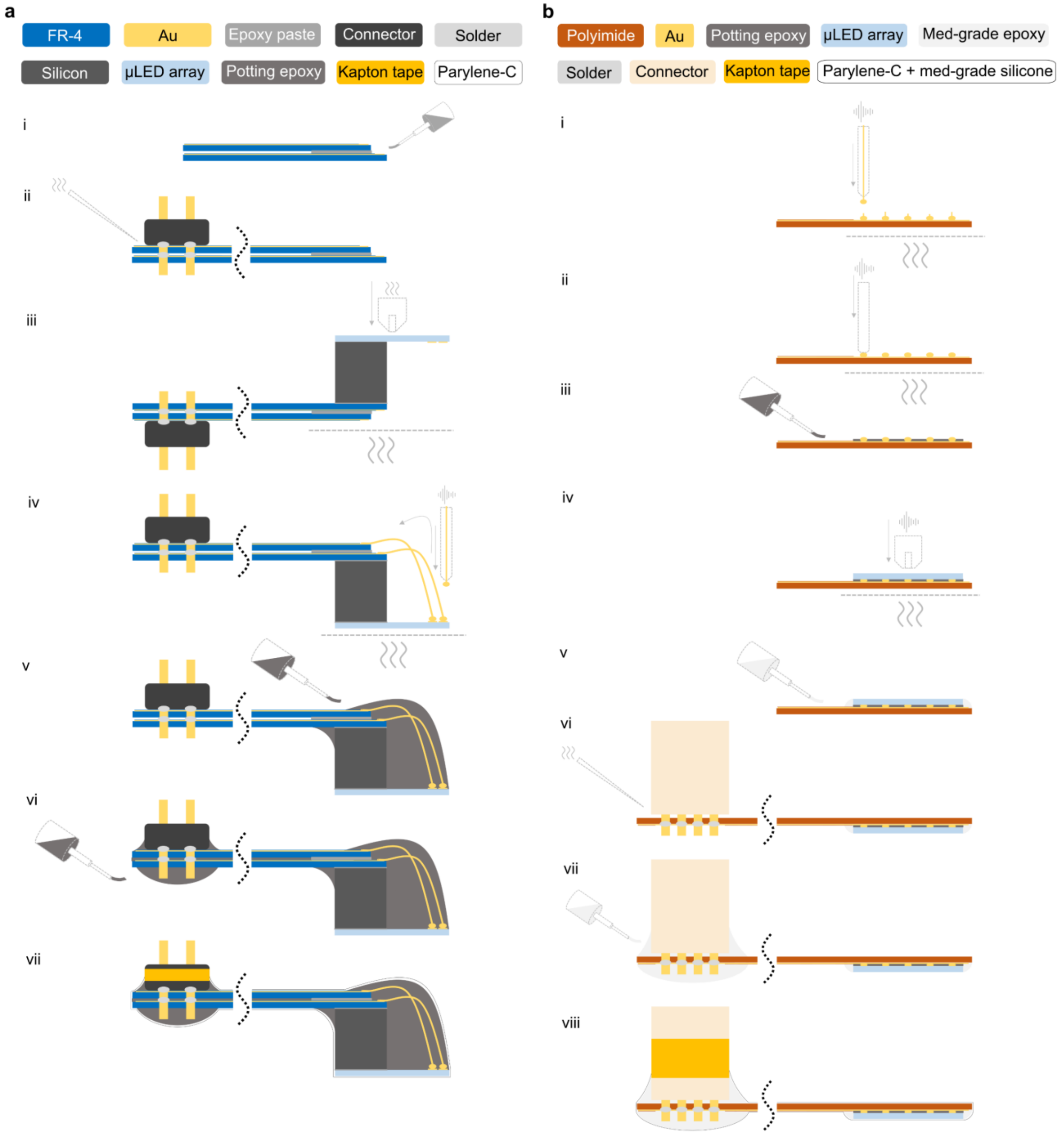


**Supplementary figure S2: Cross-sectional view of integration process for rigid (acute) and flexible (chronically implantable) systems. (a)** Bonding two 2-layer 200 µm thin FR4 PCBs (i); soldering 20-pin linear connector (ii); Bonding spacer block and µLED array using flip-chip bonding (iii); gold ball-wedge wire bonding of µLED array to PCBs (iv); potting wire bonds (v); potting connector and space between PCBs (vi); parylene-C coating, connector sealed with kapton tape before (vii). **(b)** Bump-bonding gold balls to flexible 100 µm thin polyimide PCB (i); coining bump bonds to remove excess tail (ii); underfill of bonding area prior to flip-chip bonding (iii); thermosonic flip-chip bonding of µLED array to flexible PCB (iv); medical-grade epoxy bevelled underfill around µLED array (v); soldering Omnetics connector (vi); potting Omnetics connector with medical-grade epoxy (vii); parylene-C and medical-grade silicone dispersion coating; connector sealed with kapton tape before parlyene-C coating and sapphire surface protected with UV-sensitive dicing tape prior to silicone dip coating and removed after silicone is partially cured (viii).

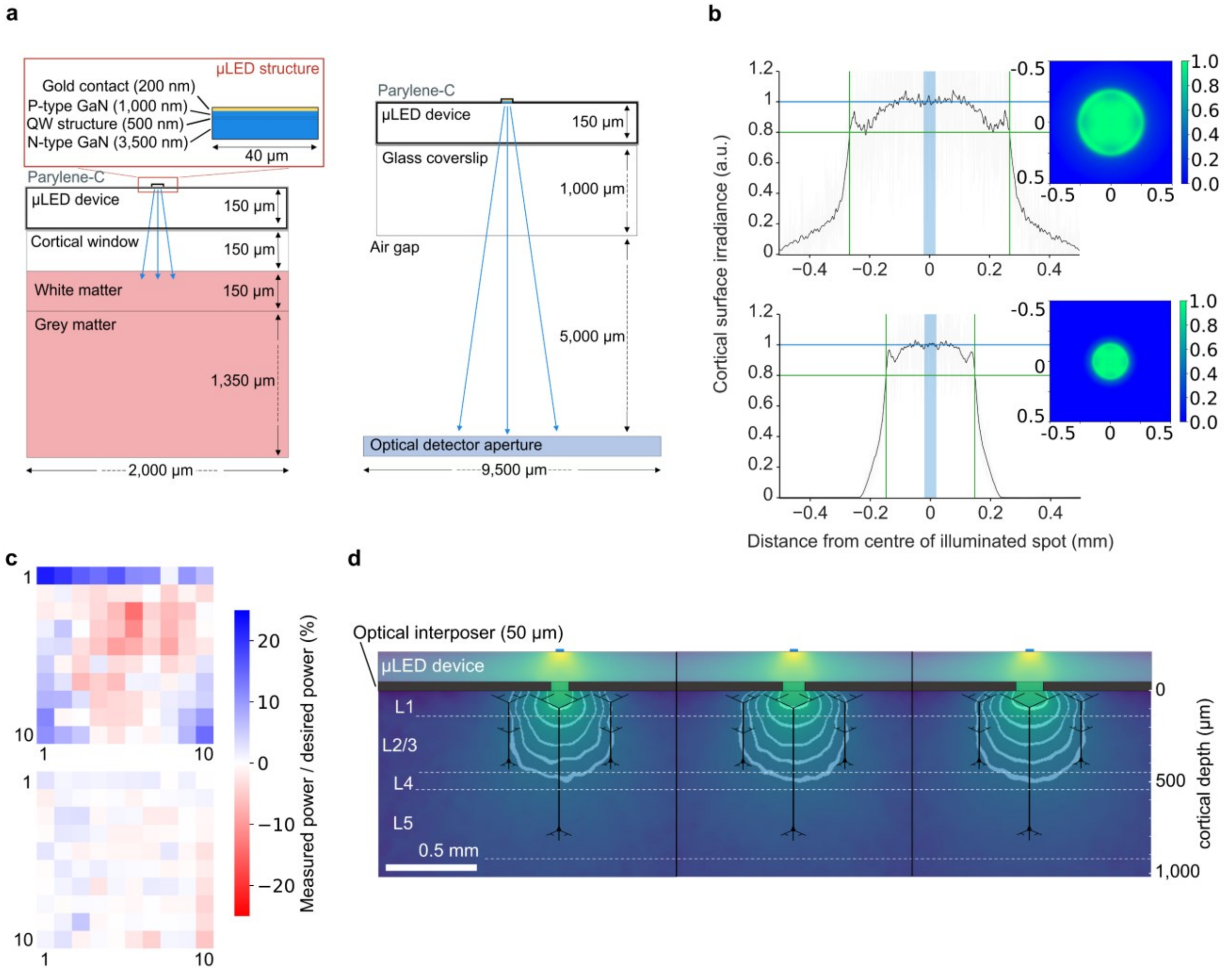


**Supplementary figure S3: Optical modelling schematics and spatial resolution enhancement through optical interposer. (a)** Schematic of optical model used to perform Monte Carlo simulations of blue light from µLED array in brain tissue (left); schematic of optical model representing laboratory setup for device calibration (right). **(b)** Central cross-section of irradiance of illuminated spot on the cortical surface with estimated spot diameter, for acute in-vivo experiment with window (top) and chronic implantation without window (bottom). Insets show full 2D irradiance map over 1x1 mm2 detector. **(c)** Normalised (measured) cortical irradiance across device for the optimal fixed drive current (top). Same but for drive currents individually inferred for each µLED from optical models (bottom). **(d)** Optical profile for different optical interposer hole diameters; left: 100 µm, middle: 125 µm, right: 150 µm. Interposer thickness is 50 µm and cortical window is not present. Grey contours show 1 mW/mm2 threshold at given depths; 100 µm, 200 µm, 300 µm, 400 µm and 500 µm. Schematic drawings of layer 5 and layer 2/3 pyramidal neurons are given for scale.

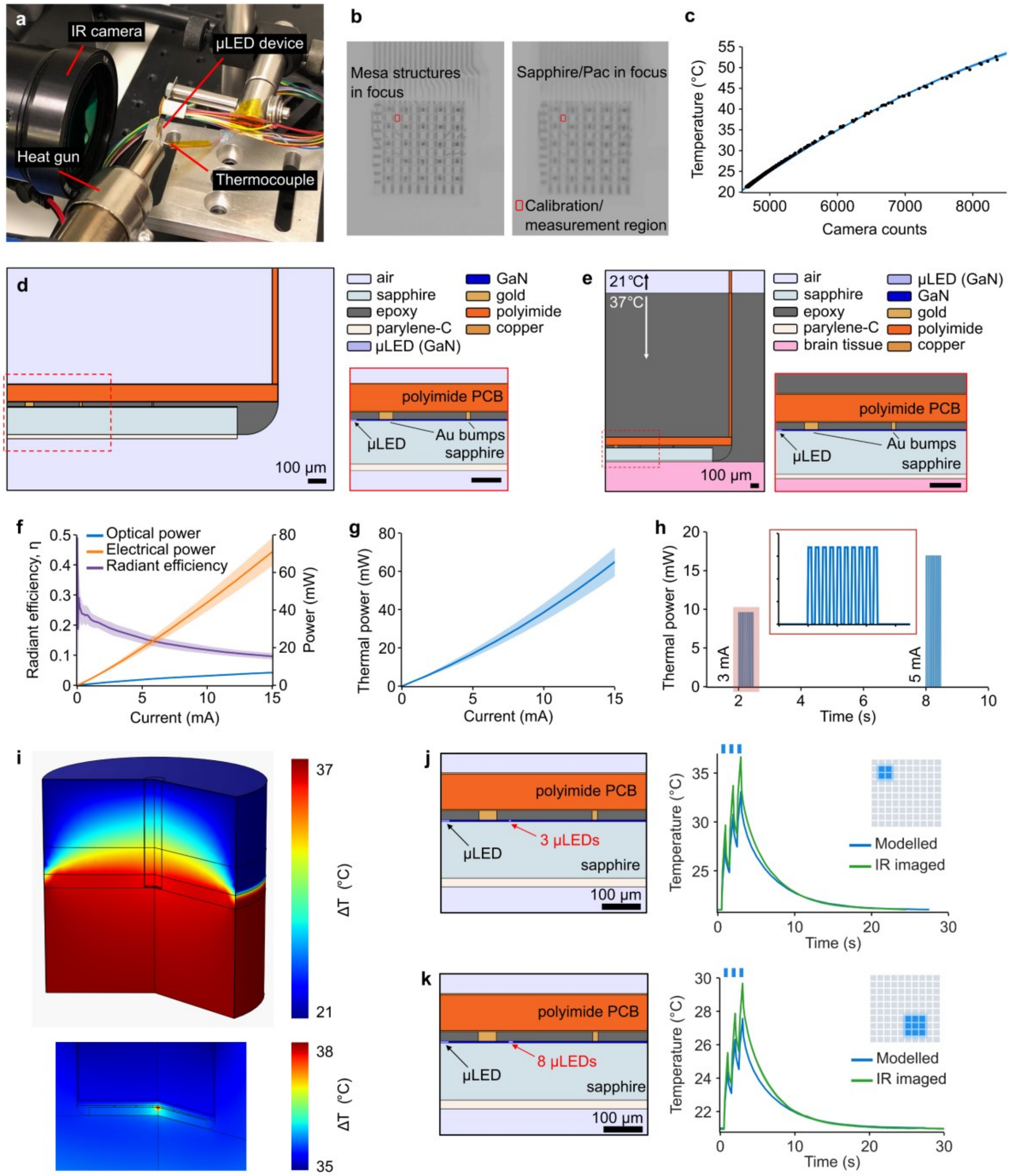


**Supplementary figure S4: Supplementary figures on thermal imaging and modelling. (a)** Photograph of thermal imaging experiment in laboratory showing infrared (IR) camera imaging sapphire surface. Thermocouple is used to measure actual temperature during calibration. **(b)** IR image showing locations used for thermal measurement and calibration, taken as an average within rectangular area. **(c)** Quadratic regression fitted to thermocouple data against camera counts; only every tenth data point is shown. **(d)** Schematic of thermal model used for direct comparison with IR imaging experiment. **(e)** Schematic of thermal model used for direct inference of thermal effects in-vivo. **(f)** Measured electrical and optical power (actual optical power generated within µLED was determined through optical models) and radiant efficiency as a function of drive current. **(g)** Thermal power as function of drive current. **(h)** Thermal pulse protocol input to thermal model for a single µLED. **(i)** Example of 3D model from COMSOL after rotation about axis of symmetry. **(j)** Comparison of IR imaging with equivalent thermal model for 2x2 square of µLEDs simultaneously illuminated; protocol is 1 Hz, 50% duty cycle, 3 mA per µLED. **(k)** Panel equivalent to (j) but for a 3x3 square of µLEDs, 1 mA per µLED.

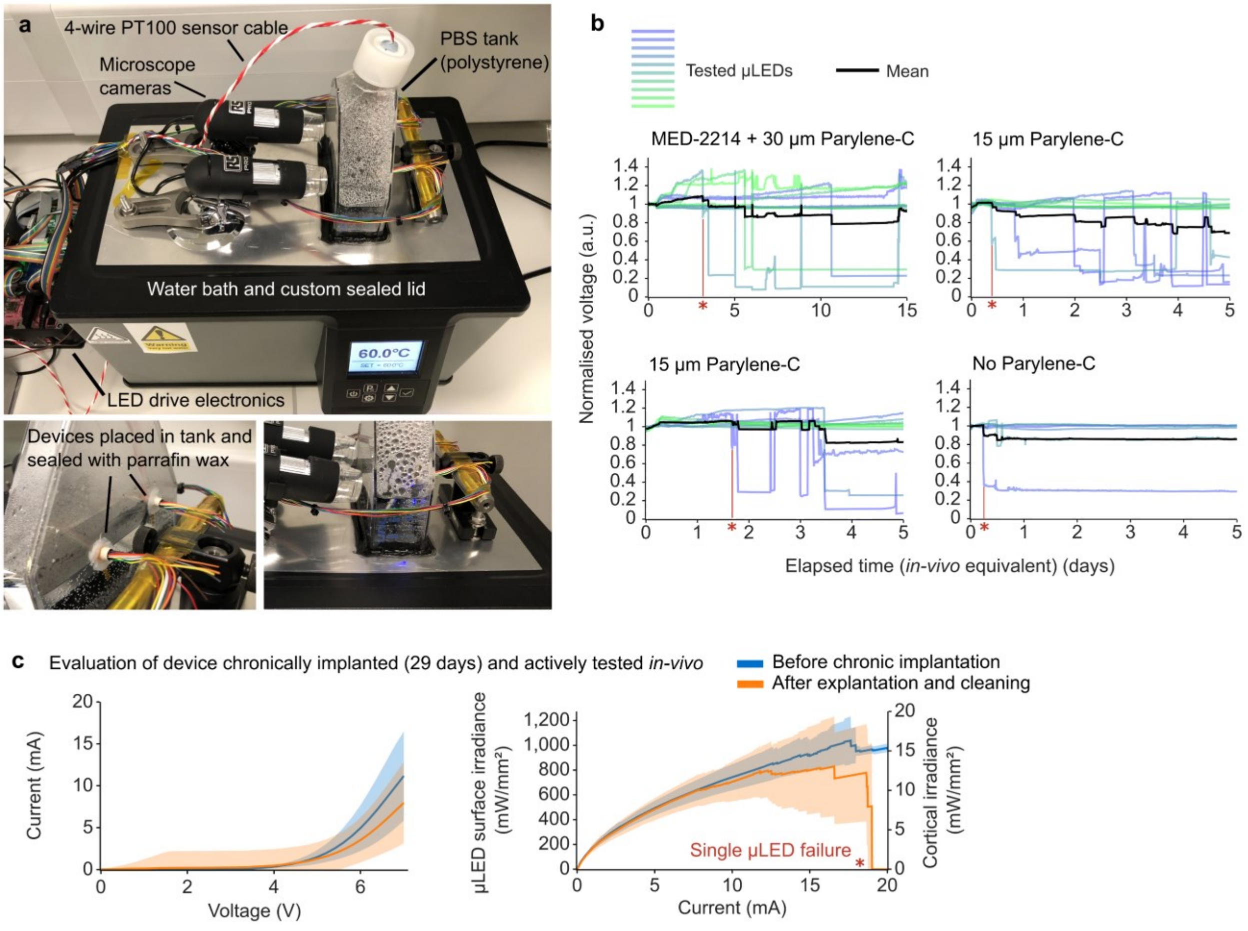


**Supplementary figure S5: Supplementary figures on accelerated ageing experiments and evaluation of chronically implanted devices. (a)** Photograph of laboratory setup showing device submerged in phosphate-buffered saline (PBS) solution heated to 60 °C using a water bath with custom lid and tank. **(b)** Normalised measured voltage during accelerated ageing for different device encapsulations; experimental protocol same as in Fig. 2d. **(c)** I-V (left) and L-I (right) before chronic implantation and after behavioural experiments and explantation of a single device.

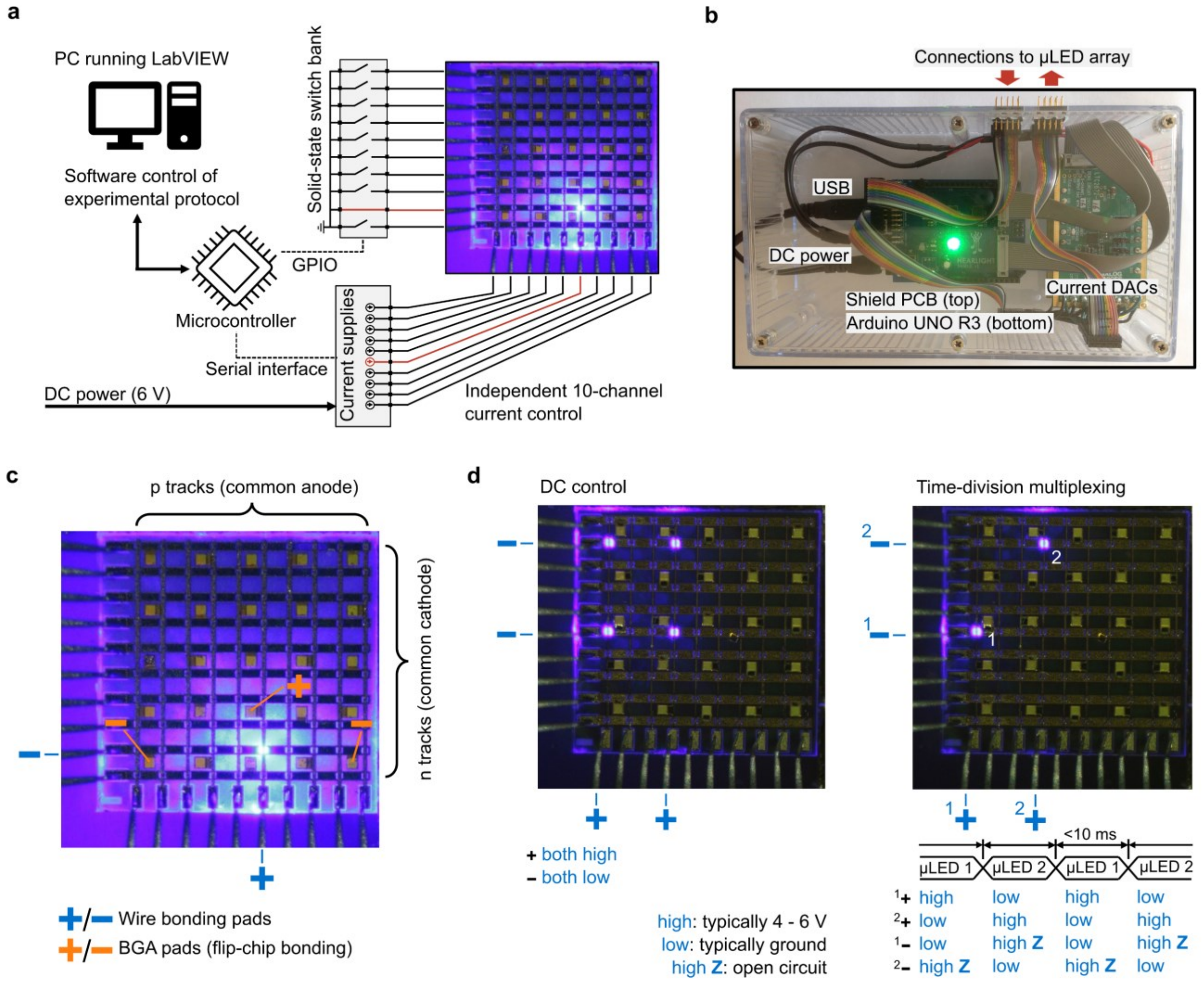


**Supplementary figure S6: Electronic controller for µLED arrays. (a)** Schematic of custom electronic control system based on current drivers operated with a microcontroller and LabVIEW interface. **(b)** Annotated photograph of the portable boxed system. **(c)** Illustration of matrix addressing of a single µLED through wire bonding or ball-grade array (BGA) pads. **(d)** Illustration of operating modes; DC which limits pattern selection (left), and time-division multiplexing allowing arbitrary patterns to be displayed, albeit at a duty cycle (right).

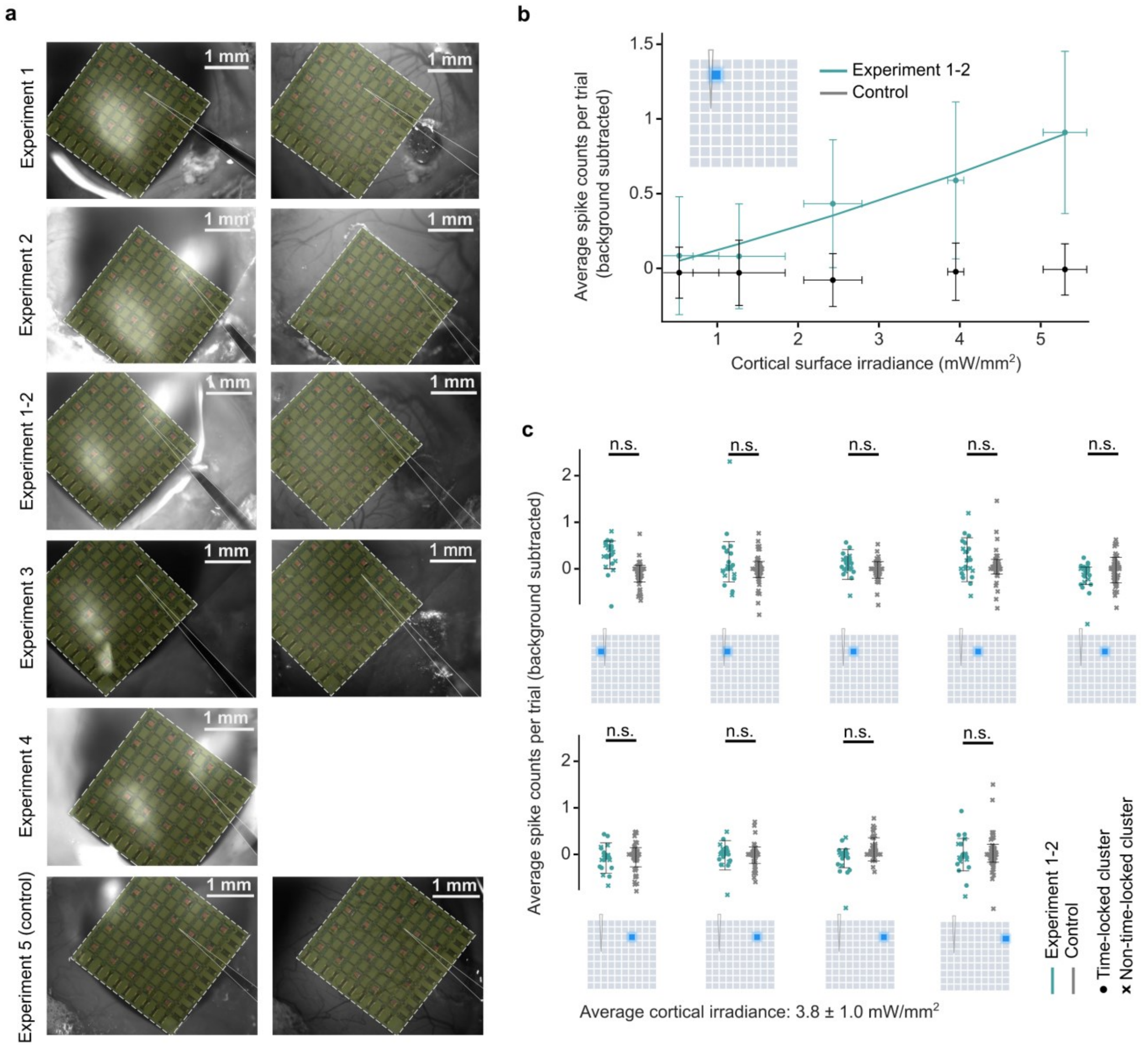


**Supplementary figure S7: Craniotomy micrographs for electrophysiology experiments and data from excluded experiment. (a)** Micrographs of craniotomy for each experiment with position of linear electrode array indicated relative to the placement of the µLED array. **(b)** Dose response for Experiment 1-2 which was omitted from main analysis due to poor recording quality (n = 12 time-locked single units for optogenetic (ChR2+) experiment; n = 51 single units for control (ChR2-) experiment). **(c)** Boxplot comparing Experiment 1-2 with control (ChR2+: n = 23, ChR2-: n = 51 single units; time-locked single units are shown as circles, non-time-locked as x's).

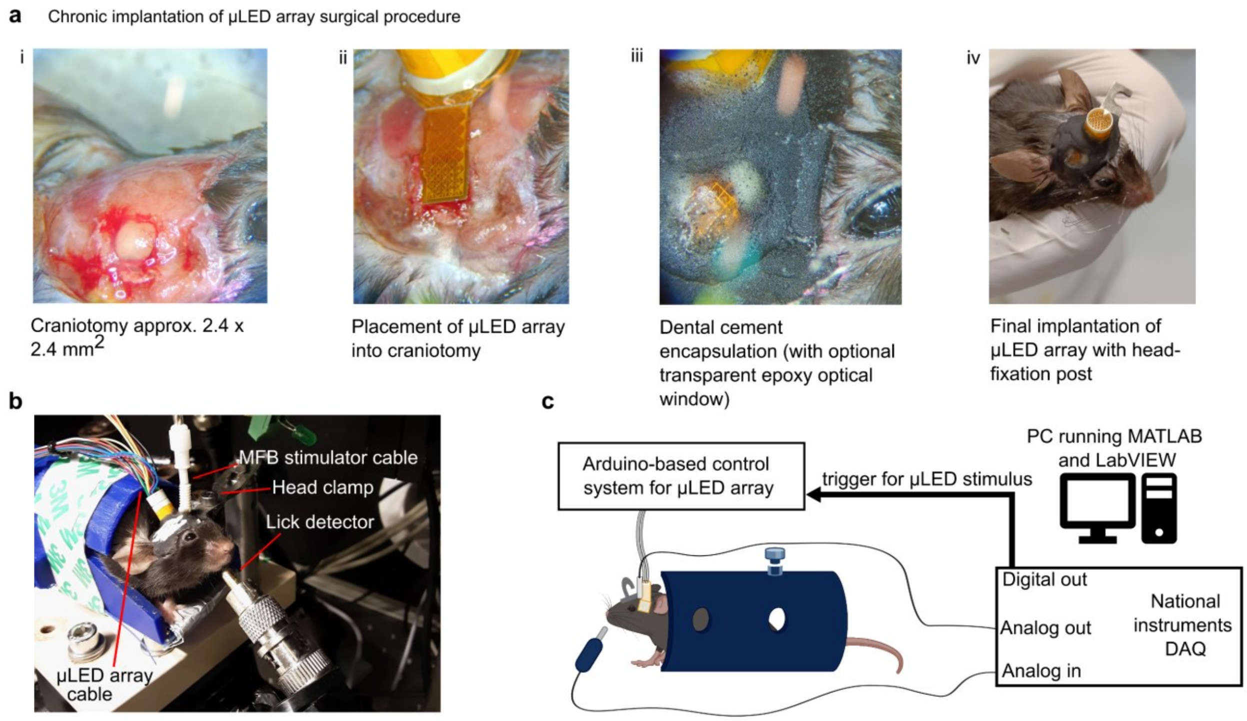


**Supplementary figure S8: Surgical steps for chronic implantation of µLED array and behavioural experimental setup.** **(a)** Photographs of surgical steps for chronically implanting the µLED array with flexible packaging. A square-shaped craniotomy is performed and the dura mater removed. The µLED array is placed into the craniotomy and secured to the skull using dental cement. **(b)** Annotated photograph of head-fixed mouse within behavioural experimental setup. **(c)** Schematic drawing of behavioural experimental setup. The experiment is managed using a MATLAB programme which specifies the protocol and decides, based on the detected lick signal, whether to apply MFB stimulation or apply time penalty. The MATLAB programme also triggers the optogenetic stimulus through the µLED control system which is separately operated using a LabVIEW programme. A National Instruments DAQ connects the lick detector, MFB stimulator and trigger signal for the µLED control system Arduino.

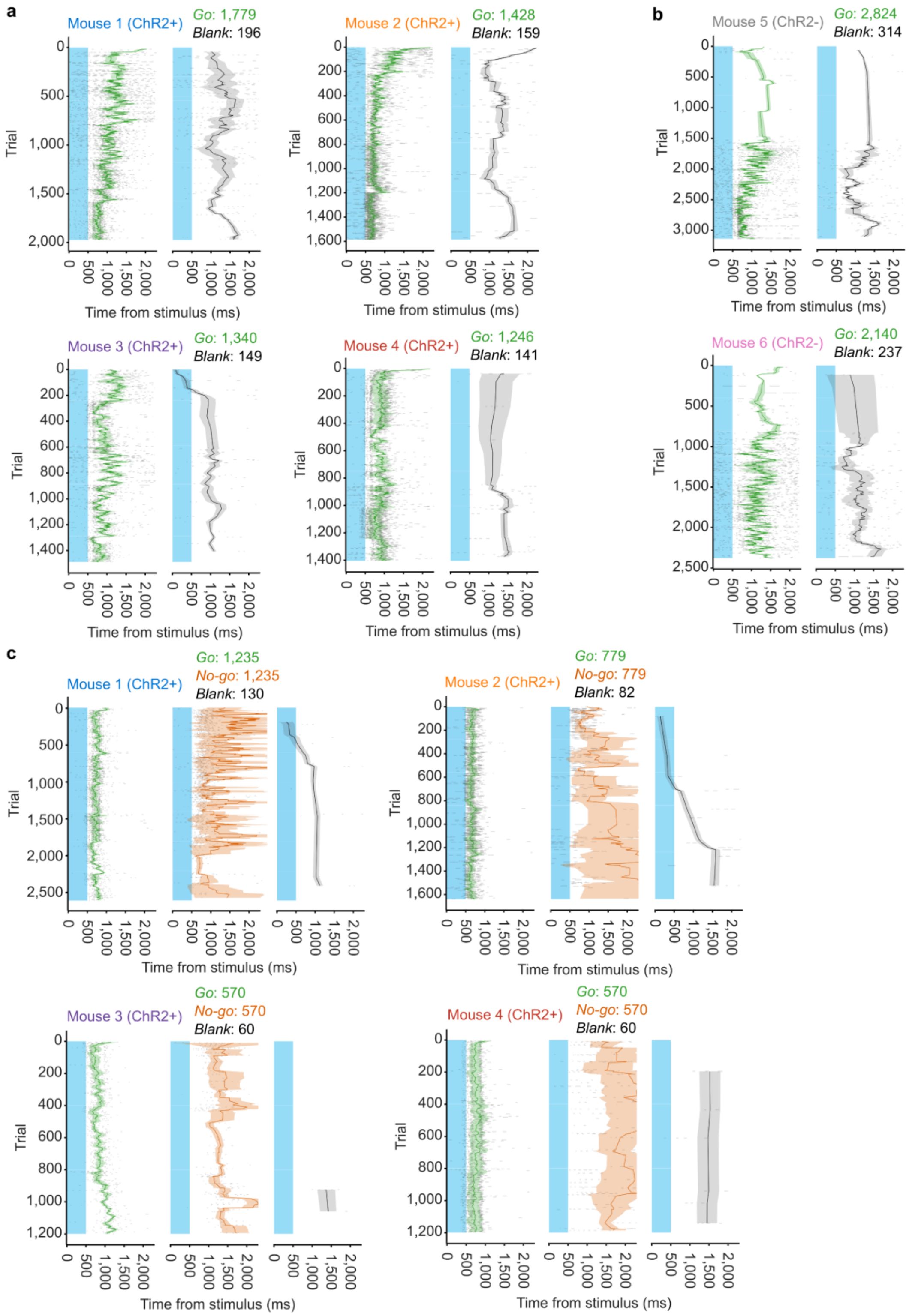


**Supplementary figure S9: Lick raster plots for each trial type for all behavioural experiments. (a)** Mean and standard deviation of lick times per trial within 2000 ms response window (10-coefficient FIR smoothing filter applied) over all sessions for the association experiment. Green: go trial licks, black: blank trial licks; light-blue shaded region shows time when µLED was illuminated in a burst of 10 pulses (20 Hz, 50% duty cycle). **(b)** Same as (a) but for the control association experiment. **(c)** Same as (a) and (b) but for the discrimination experiment. Brown-red: no-go trial licks.

| Mouse | Session # | No. trials | Accum. trials | MFB voltage (V) | MFB duration (ms) | µLED irradiance (mW/mm$^2$) | *Go* pattern |
|---|---|---|---|---|---|---|---|
| 1 | 1 | 300 | 300 | 3 | 0.01 | 10 | P8N8 P9N9 P10N10 |
| | 2 | 205 | 505 | 3 | 0.01 | 10 | P8N8 P9N9 P10N10 |
| | 3 | 233 | 775 | 3 | 0.01 | 10 | P8N8 P9N9 P10N10 |
| | 4 | 400 | 1175 | 2.4 | 0.01 | 10 | P1N1 P2N2 P3N3 |
| | 5 | 400 | 1575 | 2.4 | 0.01 | 10 | P1N1 P2N2 P3N3 |
| | 6 | 400 | 1975 | 2.4 | 0.01 | 8 | P1N4 P2N5 P3N6 |
| 2 | 1 | 187 | 187 | 1.8 | 0.01 | 10 | P1N8 P2N9 P3N10 |
| | 2 | 300 | 487 | 1.8 | 0.01 | 10 | P1N8 P2N9 P3N10 |
| | 3 | 300 | 787 | 1.7 | 0.01 | 10 | P1N8 P2N9 P3N10 |
| | 4 | 400 | 1187 | 1.7 | 0.01 | 10 | P1N8 P2N9 P3N10 |
| | 5 | 400 | 1587 | 1.7 | 0.01 | 10 | P1N8 P2N9 P3N10 |
| 3 | 1 | 239 | 239 | 2 | 0.001 | 10 | P1N1 P2N2 P3N3 |
| | 2 | 350 | 589 | 2 | 0.001 | 10 | P1N1 P2N2 P3N3 |
| | 3 | 300 | 889 | 2 | 0.001 | 10 | P1N1 P2N2 P3N3 |
| | 4 | 400 | 1289 | 2 | 0.001 | 10 | P1N1 P2N2 P3N3 |
| | 5 | 200 | 1489 | 2 | 0.001 | 6 | P3N2 P4N3 P5N4 |
| 4 | 1 | 400 | 400 | 2.6 | 0.01 | 10 | P1N1 P2N2 P3N3 |
| | 2 | 350 | 750 | 2.6 | 0.01 | 10 | P1N1 P2N2 P3N3 |
| | 3 | 255 | 1005 | 3 | 0.01 | 10 | P1N1 P2N2 P3N3 |
| | 4 | 400 | 1405 | 2.6 | 0.01 | 8 | P2N3 P3N4 P4N5 |
| 5 | 1 | 400 | 400 | 2.2 | 0.001 | 8 | P8N1 P9N2 P10N3 |
| | 2 | 400 | 800 | 2.2 | 0.001 | 8 | P8N1 P9N2 P10N3 |
| | 3 | 256 | 1056 | 2.2 | 0.001 | 8 | P8N1 P9N2 P10N3 |
| | 4 | 600 | 1656 | 2.6 | 0.001 | 8 | P8N1 P9N2 P10N3 |
| | 5 | 565 | 2221 | 2.2 | 0.001 | 8 | P8N1 P9N2 P10N3 |
| | 6 | 210 | 2431 | 2 | 0.001 | 8 | P8N1 P9N2 P10N3 |
| | 7 | 186 | 2617 | 1.9 | 0.001 | 8 | P8N1 P9N2 P10N3 |
| | 8 | 521 | 3138 | 1.7 | 0.001 | 8 | P8N1 P9N2 P10N3 |
| 6 | 1 | 400 | 400 | 1.8 | 0.01 | 6 | P1N8 P2N9 P3N10 |
| | 2 | 400 | 800 | 1.8 | 0.01 | 6 | P1N8 P2N9 P3N10 |
| | 3 | 174 | 984 | 1.8 | 0.01 | 6 | P1N8 P2N9 P3N10 |
| | 4 | 130 | 1114 | 1.8 | 0.01 | 6 | P1N8 P2N9 P3N10 |
| | 5 | 225 | 1342 | 1.9 | 0.01 | 6 | P1N8 P2N9 P3N10 |
| | 6 | 200 | 1549 | 1.6 | 0.01 | 6 | P1N8 P2N9 P3N10 |
| | 7 | 228 | 1777 | 1.2 | 0.01 | 6 | P1N8 P2N9 P3N10 |
| | 8 | 600 | 2377 | 1.5 | 0.01 | 6 | P1N8 P2N9 P3N10 |

**Supplementary table 1: Summary of behavioural association experiment parameters**

| Mouse | Session # | No. trials | Accum. trials | MFB voltage (V) | MFB duration (ms) | µLED irradiance (mW/mm$^2$) | *Go* pattern | *No-go* pattern |
|---|---|---|---|---|---|---|---|---|
| 1 | 1 | 600 | 600 | 2.4 | 0.01 | 8 | P1N4 P2N5 P3N6 | P8N8 P9N9 P10N10 |
| | 2 | 400 | 1000 | 2.4 | 0.01 | 8 | P1N4 P2N5 P3N6 | P8N8 P9N9 P10N10 |
| | 3 | 400 | 1400 | 2.3 | 0.01 | 8 | P1N4 P2N5 P3N6 | P8N8 P9N9 P10N10 |
| | 4 | 400 | 1800 | 2.3 | 0.01 | 8 | P1N4 P2N5 P3N6 | P8N8 P9N9 P10N10 |
| | 5 | 400 | 2200 | 2.2 | 0.01 | 8 | P1N4 P2N5 P3N6 | P8N8 P9N9 P10N10 |
| | 6 | 400 | 2600 | 2.2 | 0.01 | 8 | P1N4 P2N5 P3N6 | P8N8 P9N9 P10N10 |
| 2 | 1 | 280 | 280 | 1.7 | 0.01 | 10 | P1N8 P2N9 P3N10 | P5N1 P6N2 P7N3 |
| | 2 | 280 | 560 | 1.7 | 0.01 | 10 | P1N8 P2N9 P3N10 | P5N1 P6N2 P7N3 |
| | 3 | 280 | 840 | 1.7 | 0.01 | 8 | P1N8 P2N9 P3N10 | P5N1 P6N2 P7N3 |
| | 4 | 400 | 1240 | 1.7 | 0.01 | 8 | P1N8 P2N9 P3N10 | P5N1 P6N2 P7N3 |
| | 5 | 400 | 1640 | 1.7 | 0.01 | 8 | P1N8 P2N9 P3N10 | P5N1 P6N2 P7N3 |
| 3 | 1 | 400 | 400 | 2 | 0.001 | 6 | P3N2 P4N3 P5N4 | P8N8 P9N9 P10N10 |
| | 2 | 400 | 800 | 1.8 | 0.001 | 5 | P3N2 P4N3 P5N4 | P8N8 P9N9 P10N10 |
| | 3 | 400 | 1200 | 1.8 | 0.001 | 5 | P3N2 P4N3 P5N4 | P8N8 P9N9 P10N10 |
| 4 | 1 | 400 | 400 | 2.6 | 0.01 | 6 | P2N3 P3N4 P4N5 | P8N8 P9N9 P10N10 |
| | 2 | 400 | 800 | 2.6 | 0.01 | 5 | P2N3 P3N4 P4N5 | P8N8 P9N9 P10N10 |
| | 3 | 400 | 1200 | 2.6 | 0.01 | 5 | P2N3 P3N4 P4N5 | P8N8 P9N9 P10N10 |

**Supplementary table 2: Summary of behavioural discrimination experiment parameters**

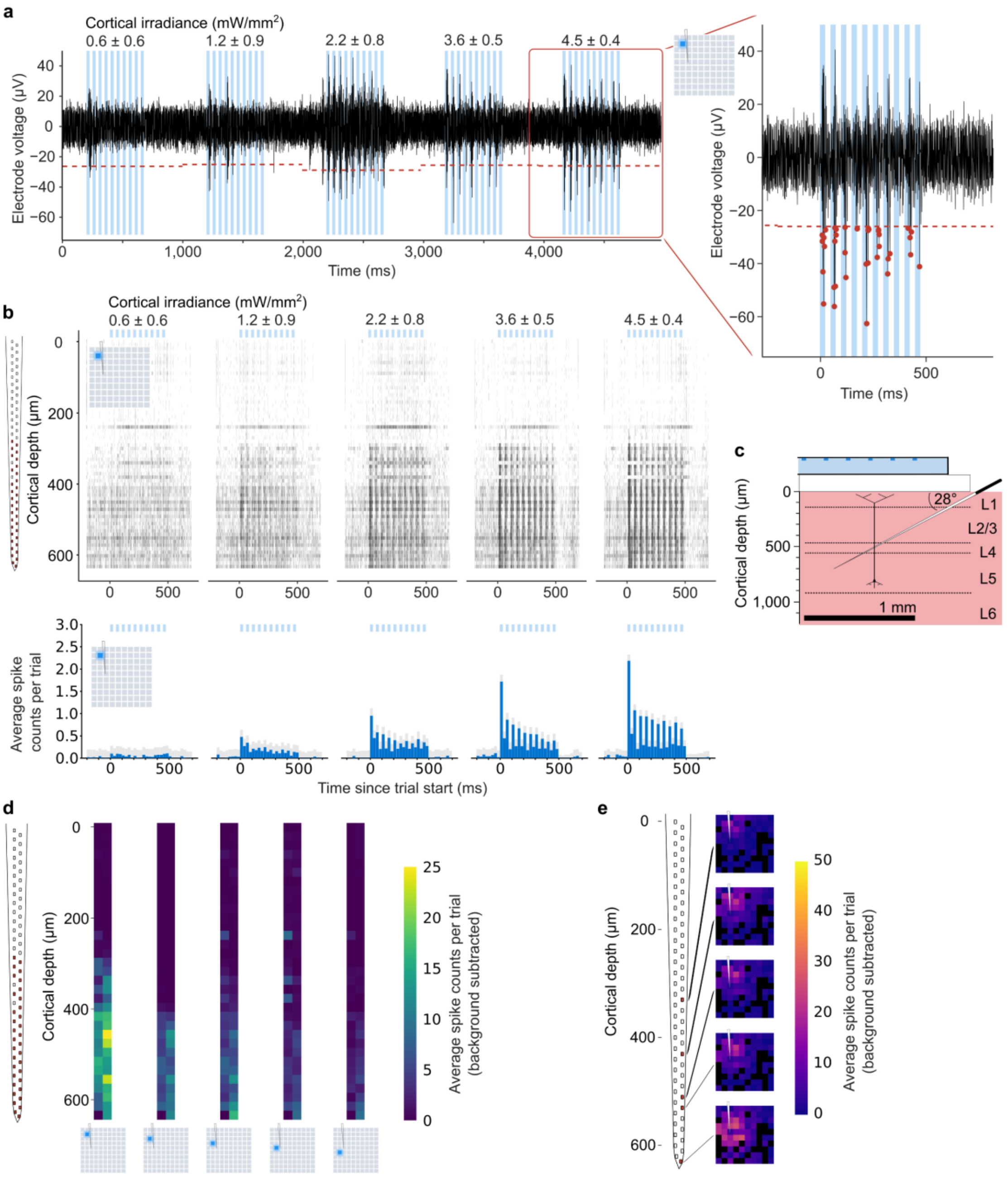


**Supplementary figure S10: Supplementary multi-unit analysis (MUA) using thresholding for Experiment 1. (a)** Example of electrophysiology recording from a single electrode (filtered using bandpass FIR filter (cutoffs 300 Hz and 5.5 kHz), common median reference subtracted, and stimulation artifact blanked using rolling average). Each µLED is pulsed 10 times at 20 Hz, 50% duty cycle for 5 cortical irradiances (mean ± standard deviation) with corresponding µLED drive currents of 2.6, 4.2, 6.1, 8.3, 10.7 mA. Right plot shows threshold (5σ from signal mean) with a "spike" identified and counted if the signal passes below and back above the threshold. **(b)** Raster plot of counted spike times under stimulation from the same µLED as (a) for each electrode on the probe, accumulated across 50 trials (top). Peri-stimulus time histogram (PSTH) plot showing binned average spike counts per trial with mean of background subtracted, averaged over all electrodes (bottom). **(c)** Schematic showing probe depth across cortical layers, for reference. **(d)** Average spike counts per trial (with background subtracted) for each electrode, as illuminated µLED position moves away from the probe. **(e)** Heat maps showing average spike counts per trial (with background subtracted) across all µLEDs individually illuminated, for selected electrode positions down the probe shank.

### Supplementary video captions

**Supplementary video 1: Patterned operation of μLED array.**

**Supplementary video 2: Behaving mouse chronically implanted with μLED array.**
